# An Interleukin 9-Zbtb18 axis promotes germinal center development of memory B cells

**DOI:** 10.1101/2023.06.11.544304

**Authors:** Xiaocui Luo, Xiaoxiao Hou, Yifeng Wang, Ye Li, Shangcheng Yu, Hai Qi

## Abstract

Germinal centers (GCs) generate humoral memory in the form of long-lived plasma cells and memory B cells (MBCs). MBC development in GCs entail profound changes in states of cell cycle, localization and survival. Whether and how these changes are extrinsically instructed and intrinsically programmed in GC B cells are not well understood. Here we demonstrate that Il-9 instructs MBC development from GCs during the primary response. Il-9 induces expression of Zbtb18, a transcription repressor that is repressed in the bulk of GC cells but highly expressed in GC memory precursor cells and MBCs. While Zbtb18 is dispensable for activation of naïve B cells and for GC formation, it is essential for normal development of GC-derived MBCs. Zbtb18 directly binds to and represses a suite of cyclin and CDK genes to promote quiescence, pro-apoptotic genes *Bid* and *Casp3* to promote survival, and GC-retaining gene *S1pr2* to promote GC departure. In the absence of Zbtb18, GCMP cells do not efficiently quit cell cycle to achieve quiescence, do not efficiently downregulate S1pr2 to exit, and they become more prone to die. Our results support that an Il-9-Zbtb18 axis instructs development of functional B-cell memory from GCs.

## Introduction

Antigen-specific memory B cells (MBCs) are of central importance to humoral immunity and rational design of antibody-based vaccines ^1^. The germinal center (GC) is a key source of MBCs. MBCs are distinct from GC B cells in three important aspects. GC B cells are actively cycling, while MBCs are largely quiescent. GC B cells are characterized by ongoing apoptosis ^2–4^, while MBCs readily survive. Unlike GC B cells that are spatially constrained and aggregated at the follicular center, MBCs are dispersed from GCs and can circulate. Therefore, MBC development from GCs must entail profound changes in these aspects in a coordinated manner. A prevailing model suggests that GCs, preferentially in the early phase of the primary response and in a manner dependent on the transcription factor Bach2, selectively export lower-affinity cells to become MBCs ^1, 5, 6^. However, while Bach2 is highly expressed in GCs and required for GC formation ^5^, Bach2 is not known to control transitional changes in those three aspects outlined above. GC-derived high-affinity MBCs exist ^7–9^, and genetic fate-mapping evidence indicates continuous MBC output from GCs throughout a primary response ^7^. Recently, homeobox protein Hhex and its interacting co-repressor Tle3 protein are found to promote MBC development, although the underlying molecular circuitry controlled by Hhex is not well understood ^10^. How MBC development from GCs is programmed, therefore, remains to be further defined.

A crucial but unsettled issue is whether MBC development from GCs are positively promoted by exogenous factors such as cytokines. We and others have previously reported that CD38^+^Fas^+^GL7^+^ GC cells contain precursors of MBCs (GCMP cells) ^9, 11, 12^. Taking advantage of a Fucci cell-cycle reporter strain, we found those precursors at the quiescent G_0_ phase can give rise to both secondary GCs and PCs upon antigen re-exposure in adoptive hosts ^9^. Follicular helper T cell-derived Interleukin 9 (Il-9) is found to promote GC exit of GCMP cells ^9^, implying an instructional model for the MBC development from GCs. On the other hand, it has been proposed that IL-9 produced by MBC themselves promotes the recall response by MBCs instead of development of MBCs ^13^.

In this study, we demonstrate IL-9 controls MBC development from GCs and identify IL-9-induced transcription repressor Zbtb18 as a key coordinator of the GC-to-MBC transitional program to promote MBC development and function.

## Results

### Il-9 promotes the development of GC-derived MBCs

Antigen-binding MBCs can develop from both GC-dependent and -independent routes, and they may function differently in the same immune response ^14–16^. To identify surrogate surface marker(s) that can distinguish GC-dependent and -independent MBCs, we resorted to the S1PR2-CreERT2×Rosa-Ai14 fate-mapping system^5^ to permanently label GC-derived cells in the model of NP-KLH immunization (Fig. 1a). We found that, irrespective of Ig isotypes, the CD73 marker ^17, 18^ can approximately label 70% GC-derived MBCs, and ∼70% of fate-mapped tdTomato^+^ MBCs were CD73^+^ (Fig. 1b-c).

**Figure 1.**
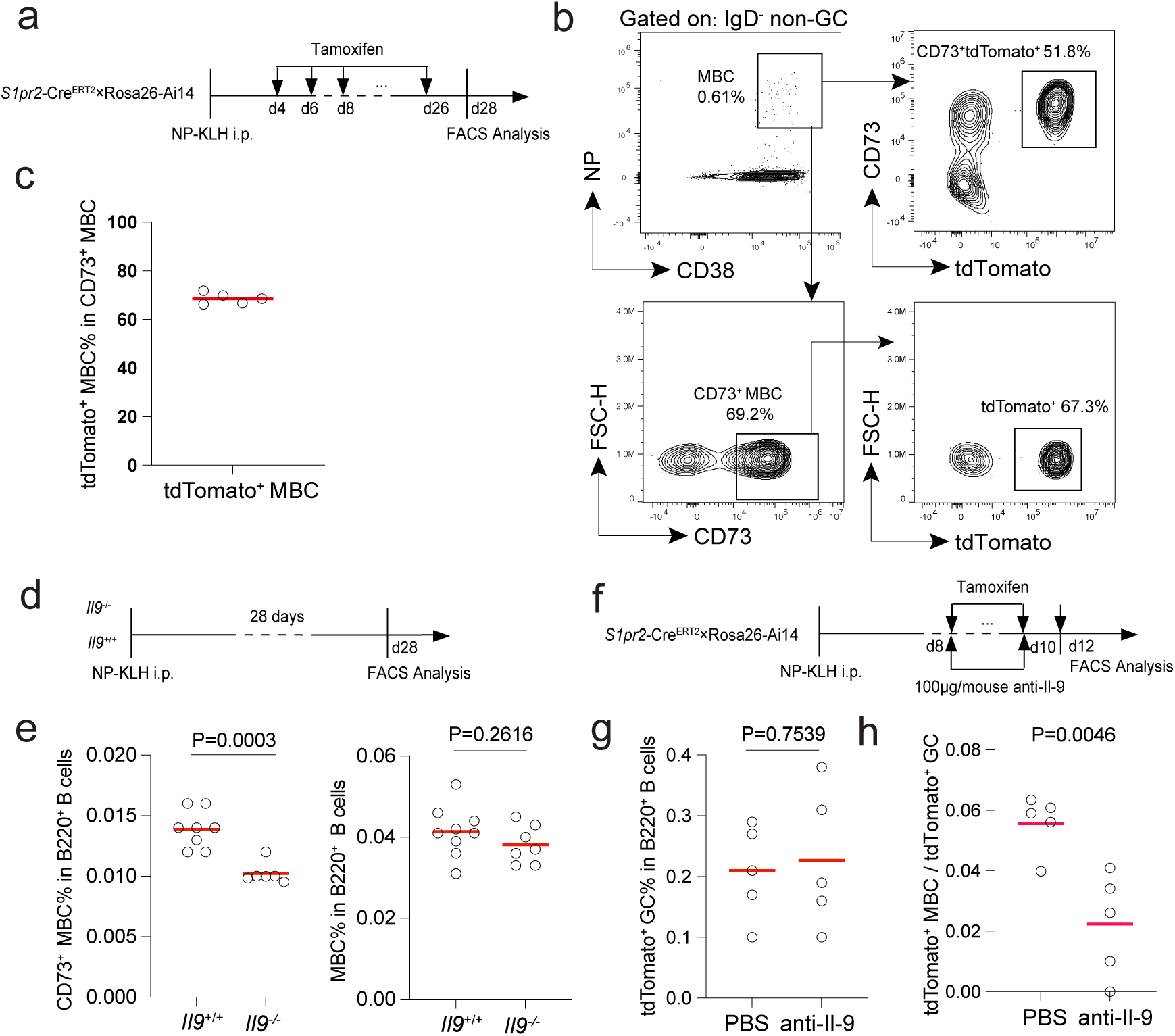
Il-9 promotes development of GC-derived MBCs. **a-c,** GC-derived MBCs in *S1pr2*-Cre^ERT2^+Rosa26-Ai14 mice 28 days after NP-KLH immunization. **a,** Experimental scheme. **b,** Representative FACS profiles and gating of tdTomato^+^ or CD73^+^ cells MBCs. **c,** Summary data of tdTomato^+^% in CD73^+^ MBCs. **d,e,** Experimental scheme (d) and summary data of CD73^+^ or total MBCs developed in *Il9*^-/-^ or *Il9*^+/+^ mice. **f,g,** Experimental scheme (f) and summary data of tdTomato^+^ MBCs as a fraction of tdTomato^+^ GCs after PBS treatment or anti-Il-9 blockade in *S1pr2*-Cre^ERT2^ξRosa26-Ai14 mice. One of three independent experiments with similar results is shown. *P* values by *t* tests.

Therefore, we used CD73 as the surrogate marker to compare abundance of GC-derived MBCs in wildtype and Il-9 knockout mice 28 days after primary NP-KLH immunization (Fig. 1d). Whereas total MBCs were largely comparable between the two types of mice, significantly fewer CD73^+^ MBCs were seen in Il-9 knockout mice (Fig. 1e). To more directly examine MBC output from GCs in a genetic fate-mapping system, we treated NP-KLH-immunized S1PR2-CreERT2×Rosa-Ai14 mice with an anti-Il-9 blocking antibody on day 8 and day 10 post immunization, while GCs were continuously labeled via tamoxifen treatment every the other day (Fig. 1f). On day 12, we enumerated tdTomato^+^ NP-binding MBCs and tdTomato^+^ NP-binding GCs and used their ratios to gauge GC output of MBCs. Il-9 blockade led to a significant reduction of MBC development from GCs (Fig. 1g-h). Taken together, these data indicate that Il-9 is required for normal development of GC-derived MBCs.

### T cell-derived Il-9 instructs MBC development

Mice deficient in the Il-9 receptor (IL9R) mount a normal primary but impaired secondary antibody response ^13^. This observation is consistent with a requirement for Il-9 in development of GC-derived MBCs. On the other hand, MBCs are proposed to produce Il-9 to enhance their own recall functions ^13^. We thus further assessed the source of Il-9 relevant for MBC development and function.

First, we immunized wildtype and IL-9-deficient mice with NP-KLH and isolated NP-binding MBCs 30 days later. An equal number of total wildtype or *Il9*^-/-^ MBCs were transferred into separate groups of ovalbumin (OVA)-primed CD45.1 mice (Fig. 2a). Seven days after NP-OVA immunization of these CD45.1 recipients, MBCs from *Il9*^-/-^ mice produced markedly fewer GCs and splenic PCs (Fig. 2b-c). These data confirm a requisite role for Il-9 in promoting B cell memory. Mechanistically, however, this role of Il-9 could be due to either that B cell-extrinsic Il-9 promotes MBC development or that B cell-intrinsic Il-9 promotes MBC recall function.

**Figure 2.**
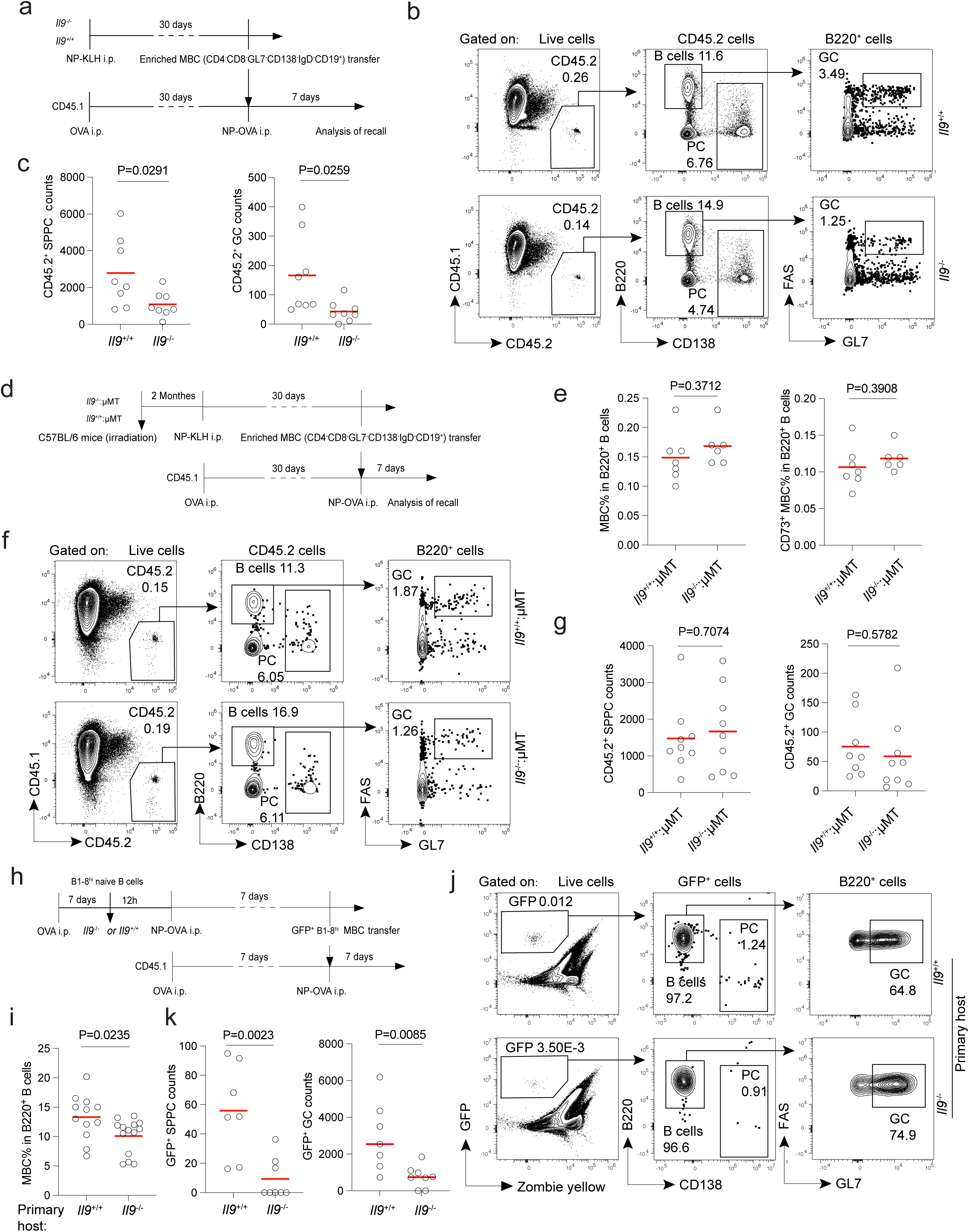
Extrinsic Il-9 is required for a functional MBC compartment. **a-c**, The recall response of MBCs isolated from *Il9*^+/+^ and *Il9*^-/-^ mice. **a,** Experimental scheme. **b,** Representative FACS profiles and gating of donor cells. **c,** Numbers of 45.2^+^ SPPC and GC cells per spleen. One of two independent experiments with similar results is shown, *P* values by *t* tests. **d-g,** The recall response of MBCs isolated from μMT:*Il9*^+/+^ and μMT:*Il9*^-/-^ chimera. **d,** Experimental scheme. **e,** Total (left) and CD73^+^ (right) MBC abundance in chimeric mice 30 days after immunization. **f,** Representative FACS profiles and gating of donor cells. **g,** Numbers of 45.2^+^ SPPC and GC cells per spleen. One of two independent experiments with similar results is shown, *P* values by *t* tests. **h-k,** The recall response of GFP-expressing B1-8^hi^ MBCs generated in *Il9*^+/+^ or *Il9*^-/-^ mice. **h,** Experimental scheme. **i,** Abundance of GFP^+^ B1-8^hi^ MBC cells generated in *Il9*^+/+^ and *Il9*^-/-^ host environment 7 days after NP-OVA immunization. Data pooled from 2 independent experiments. *P* values by *t* tests. **j,** Representative FACS profiles and gating of donor B1-8^hi^ cells in secondary hosts. **k,** Numbers of B1-8^hi^ MBC-derived SPPC and GC cells per spleen. One of three independent experiments with similar results is shown. *P* values by *t* tests.

We constructed μMT:*Il9*^-/-^ (80:20) mixed bone marrow chimera so that essentially all B cells are deficient in Il-9. Thirty days after NP-KLH immunization, these and control chimera (μMT:*Il9*^+/+^) produced comparable numbers of total and CD73^+^ NP-binding MBCs (Fig. 2d-e). We then transferred an equal number of the two types of NP-binding MBCs into ovalbumin (OVA)-primed CD45.1 mice (Fig. 2d). Seven days after NP-OVA immunization of those CD45.1 recipients, Il-9-sufficient and -deficient MBCs comparably produced secondary GCs and PCs (Fig. 2f-g). These data demonstrate Il-9 potentially produced by B cells in the primary response or during the recall of MBCs has no impact on MBC development or recall.

To further corroborate, we tested adoptive transfer of GFP-expressing B1-8^hi^ B cells into wildtype or *Il9*^-/-^ recipients. Following NP-OVA immunization, significantly fewer B1-8^hi^ MBCs were generated in *Il9*^-/-^ host animals, even though these B1-8^hi^ B cells were genetically normal (Fig. 2h-i). We then sort-purified B1-8^hi^ MBCs from these two types of host animals and transferred an equal number of such B1-8^hi^ MBCs into OVA-primed CD45.1 mice (Fig. 2h). Seven days after NP-OVA immunization of these CD45.1 recipients, wildtype B1-8^hi^ MBCs generated in *Il9*^-/-^ hosts were unable to produce comparable levels of secondary GCs and PCs in recall (Fig. 2j-k), suggesting that the MBC recall function also depends on Il-9.

Given previous findings that follicular helper T cell-derived Il-9 promotes cell cycle exit and GC departure of memory precursors ^9^, combined results from our new experiments indicate that Il-9 from T cells instead of B cells during the primary response instructs MBC development from GCs and programs MBC recall function.

### Il-9 instructs Zbtb18 upregulation in GC memory precursors

In previously published transcriptomic data ^9^, we noticed that, as compared to bulk GC B cells, CD38^+^IgD^-^Fas^+^GL7^+^ GC cells (GC memory precursors, GCMP cells) upregulated expression of *Zbtb18*. This transcription repressor, also named RP58, associates with condensed chromatins and promotes differentiation of cortical neurons in developing cerebral cortex ^19–22^, a fate-determining process that involves coordinated changes in cell cycling and positioning and is teleologically reminiscent of MBC development from GCs.

We found by quantitative RT-PCR that *Zbtb18* was expressed in naïve B cells, became markedly repressed in the bulk of Fas^+^GL7^+^ GC cells, and were upregulated in CD38^+^IgD^-^Fas^+^GL7^+^ GC cells (GC memory precursors, GCMP cells), and further increased in CD38^+^IgD^-^GL7^-^ antigen-binding MBCs (Fig. 3a). we created a knock-in reporter strain that express 3×Flag-tagged Zbtb18 from its endogenous locus (Zbtb18^Flag-in^ allele; Supplementary Fig. 1a-b). Using an anti-Flag antibody, we verified that Zbtb18 was abundantly expressed in GCMPs and MBCs, whereas it was barely detectable in the bulk of GC B cells (Fig. 3b).

**Figure 3.**
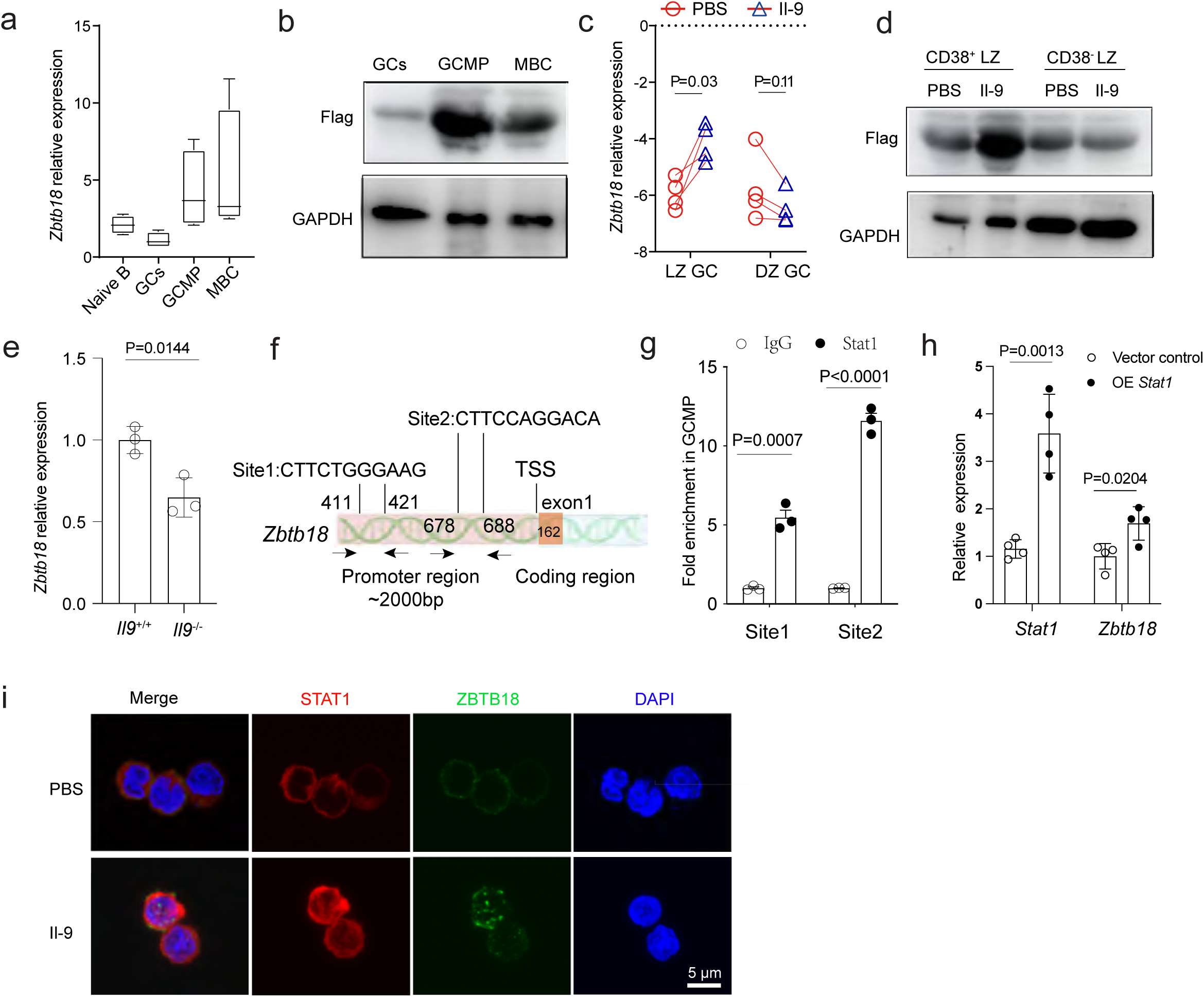
Il-9 upregulates Zbtb18 in GC memory precursors. **a,** Normalized *Zbtb18* mRNA expression in naive B cells, GCs, GCMP cells, and MBCs. Error bars indicate replicated sorts of ∼200 cells of indicated subsets from pooled splenocytes of 3 mice. One of two independent experiments with similar results is shown. **b,** Zbtb18 protein expression by GCs, GCMP cells and MBCs from *Zbtb18*^Flag-in/Flag-in^ mice. **c,** *Zbtb18* mRNA levels in light zone (LZ) or dark zone (DZ) GC B cells after *ex vivo* PBS or Il-9 treatment (200 ng/ml for 3 h). Data expressed as 1′Ct in reference to *Cd19* expression. Each line-connected pair of symbols represent one of the four independent experiments conducted. *P* values by paired *t* tests. **d,** Zbtb18 protein expression by CD38^+^ and CD38^-^ LZ GC cells of *Zbtb18*^Flag-in/Flag-in^ mice, after *ex vivo* PBS or Il-9 treatment (200 ng/ml for 3 h). One of three independent experiments with similar results is shown. **e,** The relative mRNA expression of *Zbtb18* in *Il9*^+/+^ and Il9^-/-^ GCMP cells. Data are pooled from three independent experiments. *P* values by *t* tests. **f**. Two predicted Stat1 binding sites in *Zbtb18* promoter region. Numbers indicate nucleotide positions in reference to transcription start site (TSS). Paired arrows indicate approximate positions of ChIP-qPCR primers. **g,** ChIP-qPCR for Stat1 binding to the Zbtb18 promoter in GCMP cells. Each symbol represents one independent experiment. Control IgG is set as 1. *P* values by *t* tests. **h**, Stat1 and Zbtb18 mRNA levels in B cells transduced with empty vector or Stat1-overexpressing (OE) vector, 3 days post transduction. Data represent 3 independent experiments. *P* values by *t* tests. **i,** Immunofluorescence of Stat1 and Zbtb18-Flag expression in CD38^+^ LZ GC B cells after *ex vivo* PBS or Il-9 treatment (200 ng/ml for 3 h), Scale bar: 5 μm.

Following brief Il-9 stimulation *in vitro*, *Zbtb18* expression was rapidly increased in LZ cells, particularly CD38^+^ LZ cells (essentially GCMP cells), but not in DZ GC cells (Fig. 3c-d). Conversely, GCMP cells from immunized *Il9*^-/-^ mice expressed markedly lower levels of *Zbtb18* as compared to those from wildtype mice (Fig. 3e). Il-9 in part signals through Stat1, and the *Zbtb18* promoter region contains several STAT-binding XTTCXXGGAAX motifs (Fig. 3f). By chromatin immunoprecipitation (ChIP) using an anti-Stat1 antibody, we detected significant Stat1 binding to the two sites in GCMP cells by quantitative PCR (Fig. 3g). Stat1 overexpression in B cells led to increased Zbtb18 expression (Fig. 3h). Finally, following Il-9 treatment, both Stat1 and Zbtb18 showed enhanced nuclear localization in GCMP cells (Fig. 3i). Taken together, these data indicates that Il-9 is necessary and sufficient for upregulating Zbtb18 in GCMP cells. We thus went on to explore whether Zbtb18 promotes MBC development from GCs.

### Zbtb18 is required for MBC development from GCs

We created a conditional ZBTB18 knockout allele on the B6 background (Supplementary Fig. 1c-d). We used Mb1-cre to restrict gene ablation in the B-cell lineage. Because the Mb1-cre strain was based on Cre replacement of the *Cd79a* gene, in all subsequent work we compared mice or cells of *Cd79a*^+/cre^*;Zbtb18*^fl/fl^ and *Cd79a*^+/cre^*;Zbtb18*^+/+^ genotypes, respectively termed ZBTB18^BKO^ and ZBTB18^BWT^ for convenience. We did not observe any difference in bone-marrow development of the B-cell lineage in ZBTB18^BKO^ mice (Supplementary Fig. 2). Naïve B cells from ZBTB18^BKO^ and ZBTB18^BWT^ mice were comparably activated *in vitro*, as evidenced by similar calcium mobilization in response to anti-IgM stimulation (Fig. 4a) and by similar proliferation in response to anti-IgM, LPS, or anti-CD40 stimulation (Fig. 4b-c). Following NP-KLH immunization, initial GC formation was also normal in ZBTB18^BKO^ mice (Fig. 4d-e). Therefore, Zbtb18 is dispensable for normal B-cell development, *in vitro* activation, and initial GC formation. Interestingly, at this early time of the primary response, GCMP cells as a fraction of total GCs were subtly but significantly increased, while the abundance of total NP-binding CD38^+^IgD^-^GL7^-^ MBCs in those ZBTB18^BKO^ mice showed a trend of reduction, with no difference observed in splenic PCs (Fig. 4f-h; also see Supplementary Fig. 3 for setting CD38^+^ NP-binding gates).

**Figure 4.**
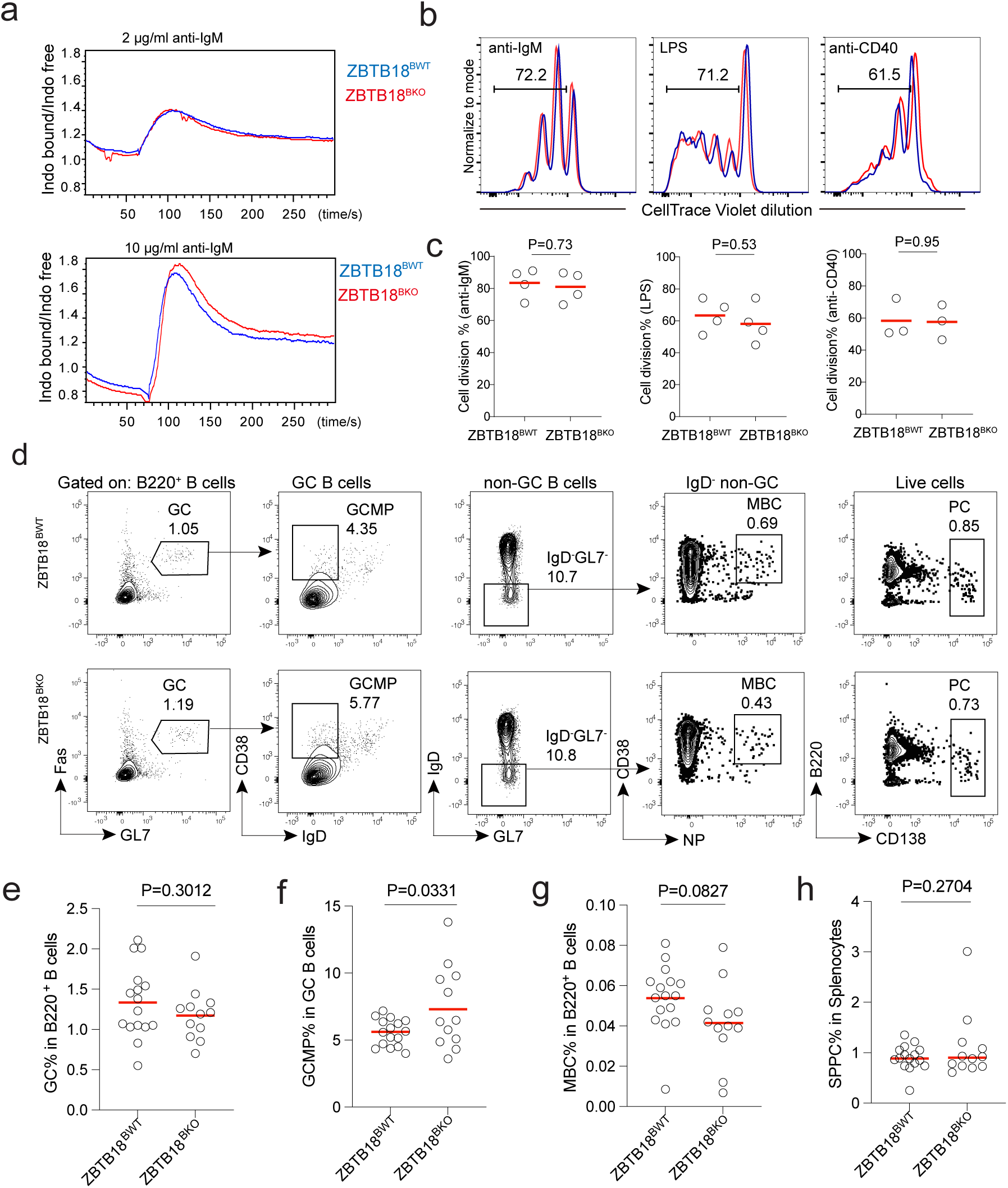
Zbtb18 does not impinge on B cell activation or GC formation. **a,** Intracellular calcium responses of naïve B cells from *Cd79a*^cre/+^ and *Cd79a*^cre/+^*Zbtb18*^fl/fl^ mice upon anti-IgM stimulation. Cells were pooled from 2-3 mice. Data represent three independent experiments. **b,c,** Proliferation of *Cd79a*^cre/+^ and *Cd79a*^cre/+^*Zbtb18*^fl/fl^ naïve B cells 72 h after anti-IgM, LPS, or anti-CD40 stimulation. Shown are representative CellTrace Violet dilution profiles (**b**) and summary data of divided fractions (**c**). Each symbol represents one independent experiment with cells pooled from 2-3 mice per genotype. *P* values by *t* tests. **d-h,** The primary immune response of *Cd79a*^cre/+^ mice and *Cd79a*^cre/+^*Zbtb18*^fl/fl^ mice 7 days after NP-KLH immunization. **d,** Representative FACS profiles and gating of GCs, GCMP cells, NP^+^ MBCs and PCs. **e-h,** Fractional abundance of GCs (**e**), GCMP cells (**f**), MBCs (**g**) and SPPCs (**h**). Data are pooled from 3 independent experiments, *P* values by *t* tests.

These observations are reminiscent of our previous findings that Il-9 blockade leads to GCMP accumulation and reduction in MBC output ^9^. We thus further examined later time points of the primary GC response, using mice that had been immunized for 14 or 28 days. At these later stages, the abundance of GCs was reduced in ZBTB18^BKO^ mice (Fig. 5a-b), whereas GCMP cells as a fraction of GCs were consistently increased, accompanied by a decrease in total MBCs and unchanged PC production (Fig. 5a, 5c-e). Because Zbtb18 is hardly expressed in the bulk of GCs but upregulated in GCMP cells (Fig. 3), these data suggest that Zbtb18 promotes MBC development from GCMP cells and export from GCs. To further corroborate, we examined *Aicda*^+/CreERT^^2^*;Zbtb18*^fl/fl^;Rosa26-Ai14 and control *Aicda*^+/CreERT^^2^*;Zbtb18*^+/+^;Rosa26-Ai14 mice that were continuously given tamoxifen to induce *Zbtb18* ablation in GCs (Fig. 5f). As shown in Fig. 5g-j, the GC abundance was comparable between the two groups, while the abundance of tdTomato^+^ GCMP cells as a fraction of tdTomato^+^ GCs was significantly increased, and the abundance of tdTomato^+^ MBCs were markedly decreased. We conclude that Zbtb18 is required for MBC development from GCs.

**Figure 5.**
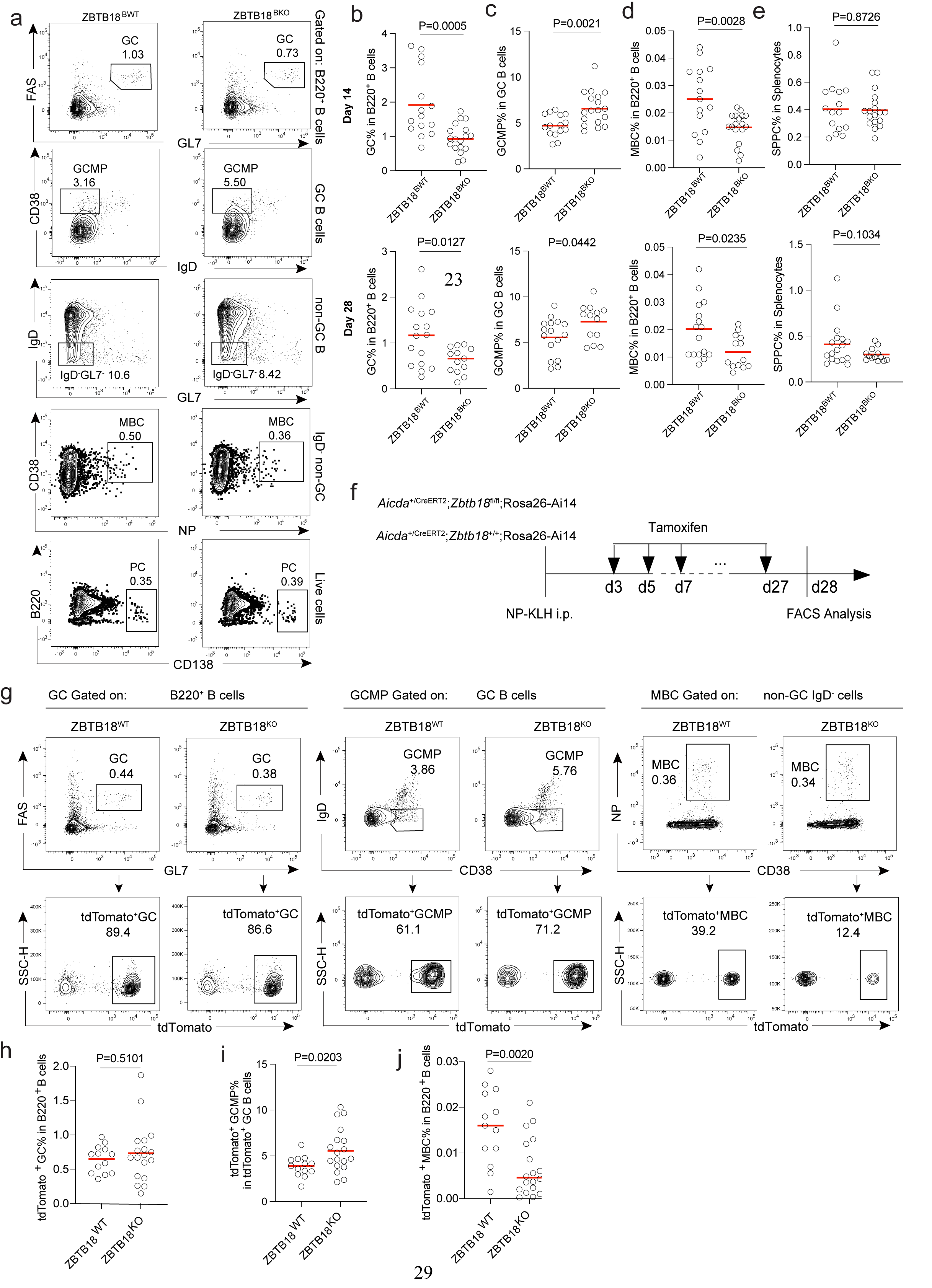
Zbtb18 is required for MBC development from GCs. **a-e,** The primary response of *Cd79a*^cre/+^ and *Cd79a*^cre/+^*Zbtb18*^fl/fl^ mice. Shown are representative FACS profiles and gating of GCs, GCMP cells, NP^+^ MBCs and SPPCs (**a**), fractional abundance of GCs (**b**), GCMP cells (**c**), MBCs (**d**) and SPPCs (**e**) 14 and 28 days after NP-KLH immunization. Data are pooled from 2 independent experiments, *P* values by *t* tests. **f-j**, The primary response of *Aicda*^+/CreERT2^;Rosa26-Ai14 and *Aicda*^+/CreERT2^;*Zbtb18*^+/+^;Rosa26-Ai14 mice. **f,** The experimental scheme. **g,** Representative FACS profiles and gating of tdTomato^+^ GCs GCMP cells, and NP^+^ MBCs. **h-j,** Fractional abundance of tdTomato^+^ GCs in total B220^+^ cells (**h**), tdTomato^+^ GCMP cells in tdTomato^+^ GCs (**i**), and tdTomato^+^ MBCs in total B220^+^ cells (**j**). Data are pooled from 3 independent experiments, *P* values by *t* tests.

### Zbtb18 programs cell cycle exit

To explore how Zbtb18 promotes MBC development, we conducted mRNA sequencing analysis of ZBTB18^BKO^ and ZBTB18^BWT^ GCMP cells. These cells differentially expressed 347 genes, including 166 upregulated and 181 downregulated in ZBTB18^BKO^ cells (*_P_*_adj_<0.05; Fig. 6a). Those expressed at higher levels in the absence of Zbtb18 were markedly enriched with genes involved in cell cycling and DNA replication (KEGG pathways, FDR *q* value<0.05; Fig. 6b), suggesting that Zbtb18 may promote MBC development by suppressing cell cycle to achieve quiescence.

**Figure 6.**
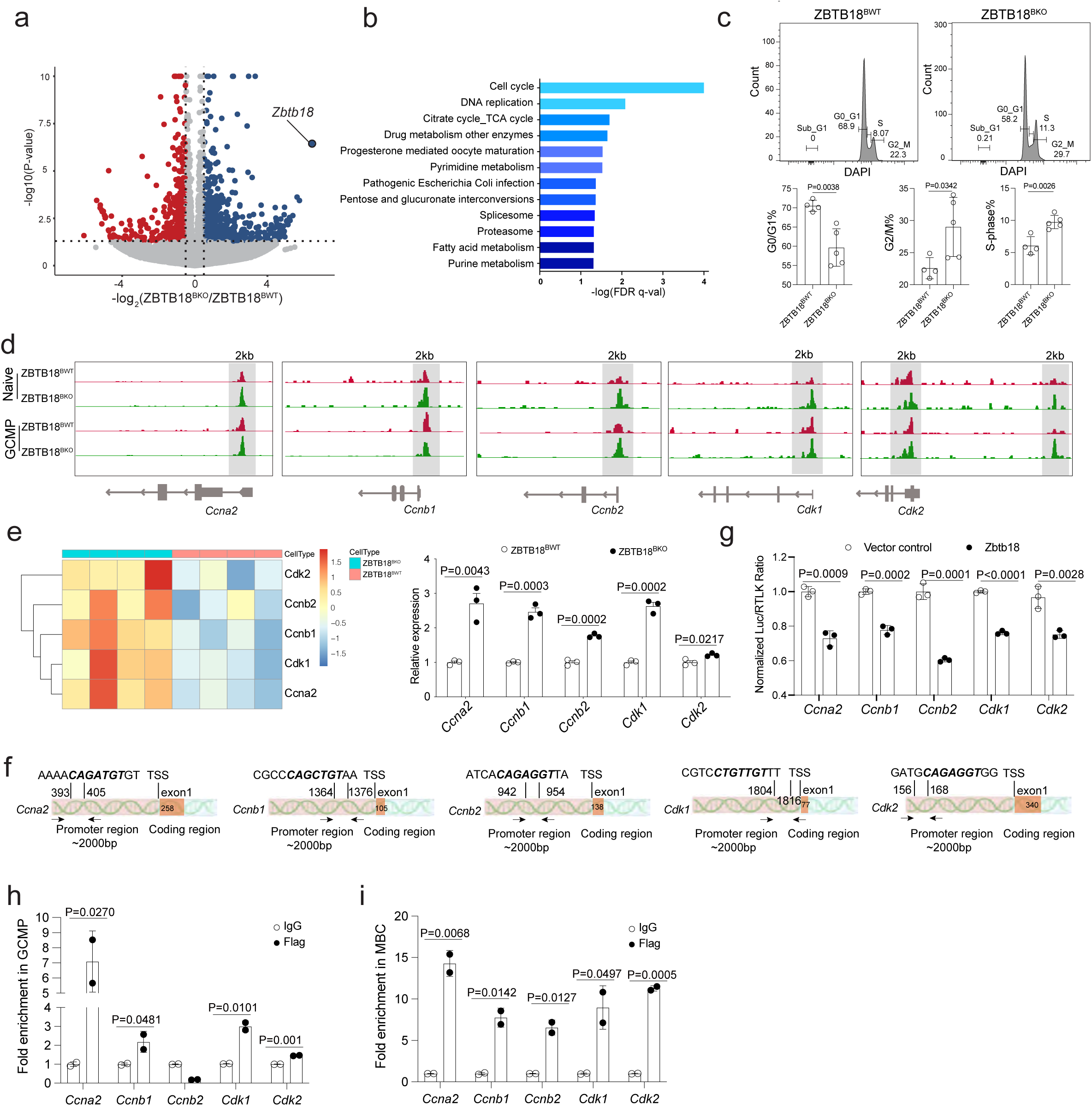
Zbtb18 promotes cell cycle exit. **a,** A volcano plot comparing transcriptomes of GCMP cells from ZBTB18^BWT^ and ZBTB18^BKO^ mice 28 days post immunization, with differentially expressed genes (*P*_adj_<0.05) highlighted and *Zbtb18* labeled. **b,** KEGG pathways enriched (FDR *q* value<0.05) in ZBTB18^BKO^ GCMP cells, based on differentially expressed genes identified in (a). **c,** Cell cycle analyses, based on DAPI-stained DNA contents, of GCMP from ZBTB18^BWT^ or ZBTB18^BKO^ mice. Flow cytometry (top) and percentage (botton) of GCMP cells in different phases are shown. Data represent one of two independent experiments. **d,** ATAC-seq read coverages around the promoters of *Ccna2, Ccnb1, Ccnb2, Cdk1* and *Cdk2* in naive B cells and GCMP cells of ZBTB18^BWT^ or ZBTB18^BKO^ mice. **e,** Expression of *Ccna2, Ccnb1, Ccnb2, Cdk1* and *Cdk2* by RNA-seq in (a), quantitated as reads count (Left), and independently examined by qRT-PCR (Right). **f-g,** Luciferase assays and ChIP-qPCR examining Zbtb18 regulation of cycle-controlling genes. **f,** Diagrams of 2-kb promoters of *Ccna2*, *Ccnb1*, *Ccnb2*, *Cdk1* and *Cdk2*, with Zbtb18-binding motifs highlighted and primers for ChIP-qPCR experiments indicated by arrows. **g,** Normalized luciferase activities, transcriptionally driven by 2-kb promoter regions of indicated genes in the presence or absence of exogenous Zbtb18. Data are from 3 independent experiments, *P* values by *t* tests. **h-i,** ChIP-qPCR analysis of interactions between Zbtb18 and indicated genes in GCMP cells (h) or MBCs (i) from *Zbtb18*^Flag-in/Flag-in^ mice. Shown are enrichment of promoter fragments of indicated genes with the anti-Flag antibody over control IgG antibody. One of two independent experiments with similar results is shown, *P* values by *t* tests.

We conducted DNA content analyses by DAPI staining and found that larger fractions of ZBTB18^BKO^ GCMP cells were in S, G2 and M phases, while a significantly smaller fraction was in the G1/G0 phase (Fig. 6c). Because Zbtb18 is a transcription repressor that is associated with heterochromatin ^19, 23^, chromatin accessibilities of Zbtb18-targeted loci are expected to increase in its absence. Therefore, we conducted ATAC-seq analysis of naïve B cells and GCMP cells of ZBTB18^BKO^ and ZBTB18^BWT^ mice. Among all annotated cell cycle-promoting genes, we obtained a list of 5 candidate genes (*Ccna2*, *Ccnb1*, *Ccnb2*, *Cdk1* and *Cdk2*) meeting two criteria: increased chromatin accessibility (Fig. 6d) and increased expression in the absence of Zbtb18 (Fig. 6e). Importantly, each of these genes contained Zbtb18 consensus binding motifs within the 2-kb region upstream of their transcription start sites (Fig. 6f). We cloned the 2-kb promoter region from these 5 genes to conduct luciferase assays and found that Zbtb18 repressed promoter activities of all these 5 genes (Fig. 6g). Using primer sets surrounding Zbtb18 consensus binding motifs, we detected significant Zbtb18 binding to *Ccna2*, *Ccnb1*, *Cdk1* and *Cdk2* by ChIP-qPCR in GCMP cells isolated from immunized *Zbtb18*^Flag-in/+^ mice (Fig. 6f and 6h). In addition, significant Zbtb18 binding to all 5 genes was detected in MBCs isolated from immunized *Zbtb18*^Flag-in/+^ mice (Fig. 6i).

Therefore, Zbtb18 directly represses critical cell cycle-promoting genes and promotes cell cycle exit, which would in turn favor the GC-to-MBC transition.

### Zbtb18 promotes survival

In the DNA content analysis of ZBTB18^BKO^ GCMP cells, we noticed a subtle but consistent increase of sub-G1 events (Fig. 6c), suggestive of increased apoptosis. Supporting this possibility, a markedly increased fraction of ZBTB18^BKO^ GCMP cells contained active caspases (Fig. 7a). Among known apoptosis-promoting genes, *Casp3* and *Bid* were expressed at higher levels in ZBTB18^BKO^ GCMP cells (Fig. 7b). Promoter regions of these two genes contained Zbtb18 consensus binding motifs, and their promoter activities are suppressed by Zbtb18, as measured by luciferase assays (Fig. 7c). In MBCs isolated from immunized *Zbtb18*^Flag-in/+^ mice, we confirmed Zbtb18 binding to promoters of *Casp3* and *Bid* (Fig. 7d). Together, these data indicate that Zbtb18 directly represses *Casp3* and *Bid* expression and promotes survival of GCMP cells.

**Figure 7.**
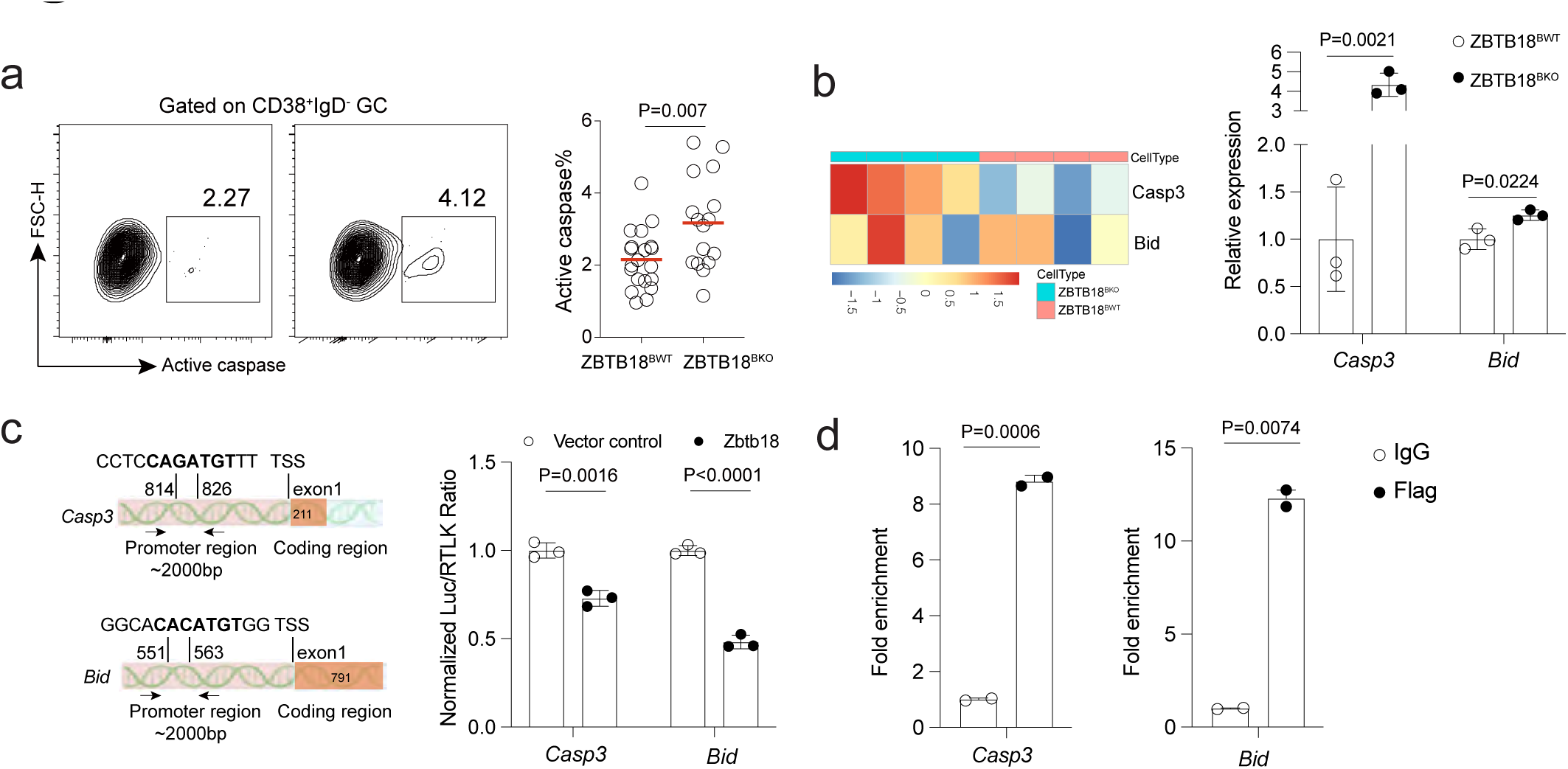
Zbtb18 promotes survival of GCMP cells. **a,** Representative FACS profiles and summary data of GCMP cells expressing active caspases. In scatter plots, each dot represents one mouse, and lines demote means. Data are pooled from four independent experiments. *P* values by *t* tests. **b,** Expression of *Casp3* and *Bid* by RNA-seq in Fig. 6a, quantitated as reads count (Left), and independently examined by qRT-PCR (Right). **c,** Left, diagrams of 2-kb promoters of *Casp3* and *Bid* with Zbtb18-binding motifs highlighted and primers for ChIP-qPCR experiments indicated by arrows. Right, normalized luciferase activities, transcriptionally driven by 2-kb promoter regions of indicated genes in the presence or absence of exogenous Zbtb18. One of two independent experiments with similar results is shown, *P* values by *t* tests. **d,** ChIP-qPCR analysis of interactions between Zbtb18 and indicated genes in MBCs from *Zbtb18*^Flag-in/Flag-in^ mice. Shown are enrichment of promoter fragments of indicated genes with the anti-Flag antibody over control IgG antibody. One of two independent experiments with similar results is shown. *P* values by *t* tests

### Zbtb18 programs GC departure

In the absence of Zbtb18, the fractional abundance of GCMP cells in GCs was increased and the abundance of MBCs in the spleen was decreased (Fig. 4 and Fig. 5), similar to what was previously observed when IL-9 was blocked ^9^. These data indicate that GC exit, another prerequisite for MBC development, depends on a Zbtb18-controlled pathway.

S1pr2 is a Gα12/13-coupled chemo-repulsive receptor that is highly expressed by GC B cells and obligately maintain the dense packing of GC cells in the follicular center. S1PR2 must be downregulated as cells leave GCs ^9, 11, 24^. By quantitative RT-PCR we found that ZBTB18^BKO^ GCMP cells more strongly expressed *S1pr2* than ZBTB18^BWT^ counterparts (Fig. 8a). Furthermore, we conducted a transwell assay in which S1P-mediated inhibition of GCMP migration was tested in conjunction with the S1pr2-specific antagonist JTE-013. As shown in Fig. 8b, S1P achieved stronger inhibition of ZBTB18^BKO^ than ZBTB18^BWT^ cells, and such difference was abrogated by JTE-013. These data indicate that ZBTB18^BKO^ cells expressed higher levels of functional S1pr2 receptor. A Zbtb18-binding motif was found 30 bp upstream of the *S1pr2* transcription starting site (Fig. 8c). By luciferase assays, we found that a 2-kb promoter region conferred Zbtb18-depedent repression (Fig. 8d). In GCMP cells isolated from immunized *Zbtb18*^Flag-in/+^ mice, we verified direct binding of Zbtb18 to the *S1pr2* promoter by ChIP-qPCR (Fig. 8e). Finally, in the absence of Zbtb18, GCMP cells were positioned further away from the edge of GCs, indicative of retarded exit (Fig. 8f-g). Taken together, these data indicate that Zbtb18 directly represses *S1pr2* in GCMP cells and programs their exit from the GC.

**Figure 8.**
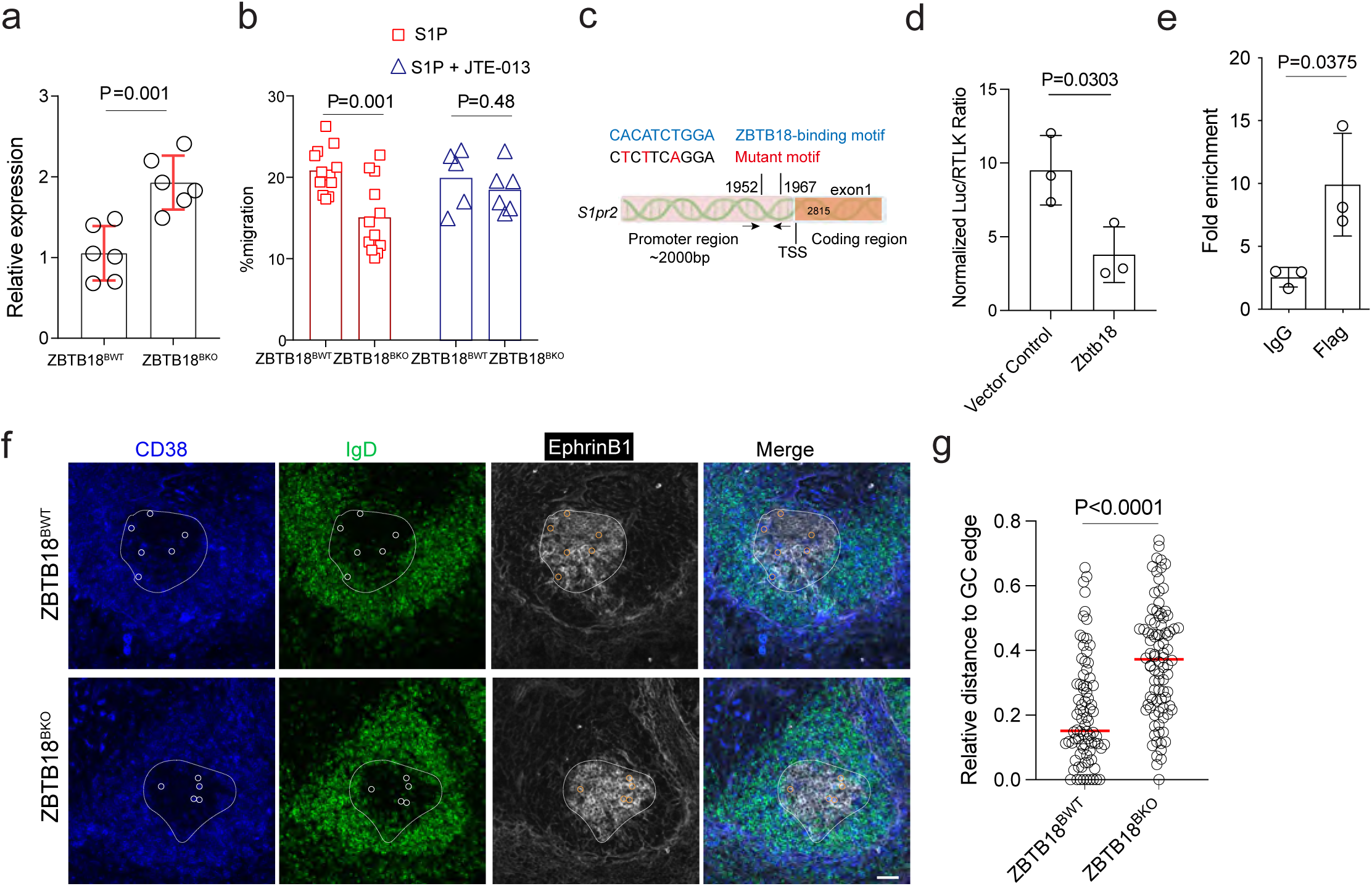
Zbtb18 suppresses *S1pr2* to program GC departure. **a,** *S1pr2* expression in GCMP cells of indicated genotypes by qRT-PCR. Each dot represents an independent sort of 200 cells. One of two independent experiments is shown. *P* value by *t* test. **b,** Transwell migration of GCMP cells subjected to inhibition by S1P or S1P plus S1PR2 antagonist JTE-013. Each dot represents one transwell, and data are pooled from two independent experiments. *P* values from *t* tests. **c,** Diagram of the 2-kb *S1pr2* promoter, with position of Zbtb18-binding motifs indicated. Primers for ChIP-qPCR experiments indicated by arrows. **d,** Normalized luciferase activities transcriptionally driven by a 2-kb *S1pr2* promoter containing the Zbtb18-binding motif, in the presence or absence of exogenous Zbtb18. Three independent experiments are shown. *P* values by paired *t* tests. **e,** ChIP-qPCR analysis of interactions between Zbtb18 and the *S1pr2* promoter in *Zbtb18*^Flag-in/Flag-in^ cells. One of two independent experiments with similar results is shown. **f**, Tissue distribution patterns of GCMP cells in ZBTB18^BWT^ or ZBTB18^BKO^ GCs. Demarcation of GC border (white circle) were based on IgD and Ephrin B1 staining. Orange circles: CD38^+^IgD^-^Efnb1^+^. Scale bar: 40 μm. **g**, Relative distance to the GC edge, calculated as the shortest distance to the GC edge divided by the GC radius. More than 7 GCs from 3 mice in each group were analyzed. *P* value by t test.

### MBC development and function controlled by an Il-9-Zbtb18 axis

Finally, we re-examined Zbtb18-controlled genes in GCMP cells from *Il9*^-/-^ mice. Compared to the control, GCMP cells developing in the absence of Il-9 expressed abnormally high levels of *Ccna2*, *Ccnb1*, *Ccnb2*, *Cdk1*, *Cdk2*, *Casp3*, *Bid*, and *S1pr2* (Supplementary Fig. 4a). Conversely, Il-9 treatment of GCMP cells *in vitro* led to downregulation of a majority of these genes (Supplementary Fig. 4b), an effect that was significantly attenuated when Zbtb18 was absent (Supplementary Fig. 4c). Finally, impaired MBC formation in *Il9*^-/-^ mice can be largely rescued when Zbtb18 expression was enforced in antigen-specific B cells by retroviral transduction (Supplementary Fig. 4d-e). Combined together, our results support that an Il-9-Zbtb18 axis promotes MBC development and function.

## Discussion

Our study demonstrates that Il-9 instructs MBC development from GCs. By analyzing behaviors of MBC precursors in GCs, we have previously suggested that Il-9 from T cells is required for normal MBC development from GCs ^9^. However, that early work did not specifically quantitate GC-derived MBCs or examine their recall. By analyzing serum antibodies over time in *Il9r*^-/-^ mice, Takatsuka and colleagues propose that Il-9 signaling to B cells is important for recall antibody responses ^13^. While the defect in recall antibody titers could be explained by defective formation of GC-derived MBCs during the primary response in the absence of Il-9 signaling to B cells, those authors instead suggest that Il-9 derived from MBCs themselves promote their own recall functions, as they did not observe defective IgG1 MBC development in the absence of Il-9 signaling to B cells. However, quantitation of IgG1 MBCs at the end of the primary response does not take into account of MBCs of IgM and other switched isotypes and could not accurately account for all GC-derived MBCs.

By series of chimera and adoptive transfer experiments and by quantifying all GC-derived MBCs specifically, we have now provided definitive evidence that Il-9 from T cells but not B cells promotes MBC development from GCs. Moreover, by separating hosts of the primary and secondary responses, we demonstrate that genetically normal MBCs generated in Il-9-deficient primary hosts cannot be efficiently recalled in secondary hosts that are sufficient for Il-9, further highlighting an essential role for Il-9 in programming B cell memory during the primary response.

Another key advance of the current study is the revelation that Il-9 induces Zbtb18 in GCMP cells, and Zbtb18 in turn promotes GCMP cells to stop cycling, acquire an improved ability to survive, and exit GCs to become MBCs. Consistent with these functional effects, Zbtb18 directly represses a suite of genes that promote cell cycling (e.g. *Ccna2*, *Ccnb1*, *Ccnb2*, *Cdk1* and *Cdk2*), potentiate apoptosis (e.g. *Casp3* and *Bid*), and maintain GC retention (e.g. *S1pr2*). Retroviral Zbtb18 overexpression in B cells leads to repression of multiple genes in the PI3K pathway ^25^. Consistent with those observations, chromatin accessibilities of multiple PI3K pathway-related genes were increased in ZBTB18^BKO^ cells according to our ATAC-seq analyses (data not shown). Activities of the PI3K pathway are positively linked to mTORC1 activities, which in GCs promote biomass accumulation and proliferation after cyclic re-entry ^26^. We speculate that Zbtb18 could promote MBC development by this additional mechanism of dampening PI3K and mTORC1 activities to inhibit DZ re-entry.

Transcription factor Hhex has been reported to promote MBC development ^10^. Similar to Zbtb18, Hhex is repressed in GCs and upregulated in GCMP cells. Unlike Hhex and Zbtb18, Bach2 is expressed highly in the bulk of GCs and not upregulated in GCMP cells ^9, 10^. In cooperation with co-repressor Tle3, Hhex is suggested to repress *Bcl6* and thereby release Bcl-6-mediated suppression of Bcl-2. When the *Hhex* gene is ablated at the GC stage, the fraction of GCMP cells is reduced in GCs, and MBCs are reduced. When *Zbtb18* is deleted at the GC stage, the fraction of GCMP cells is increased in GCs, partly because of retarded GC exit, and MBCs are reduced. Therefore, a plausible scenario is that Hhex and Zbtb18 function sequentially, with the former acting to enhance the general survivability of GCMP cells and the latter acting to promote cycle arrest, further survivability increase and GC departure. In this context, it is interesting to note that Hhex expression in LZ GC cells is upregulated by Il-9 (data not shown). The possibility that Hhex and Zbtb18 act in tandem downstream of Il-9 to promote MBC development is consistent with our observation that the defect resulting from an Il-9 deficiency tends to be more severe than a Zbtb18 deficiency.

In summary, by demonstrating Il-9 as a pre-requisite for normal development of GC-derived MBCs and by identifying Il-9-controlled Zbtb18 as a key transcription factor that coordinates GC-to-MBC transition, our work support an instructional model for the development of GC-derived memory.

## Acknowledgements

We thank X. Liu, B. Liu and T. Shao for critical comments of the manuscript. We thank M. Nussenzweig, M. Reth and T. Kurosaki for sharing mice. The work was funded in part by the National Key R&D Program of China (Ministry of Science and Technology, 2018YFE0200300 to H.Q.), National Natural Science Foundation of China (grant 31830023 and 81621002 to H.Q.), the Changping Laboratory, the Tsinghua-Peking Center for Life Sciences, the Beijing Municipal Science & Technology Commission, and the Beijing Frontier Research Center for Biological Structure. H.Q. is a New Cornerstone Investigator.

## Author contributions

H.Q. conceptualized and supervised the study. X.L. and X.H. conducted a majority of the experiments together with Y.W. Y.L. helped with RNA-seq and ATAC-seq analysis. S.Y. conducted ELISA experiments. H.Q. and X.L. wrote the paper with input from all authors.

## Data availability

All data generated and/or analyzed during the current study are available from the corresponding author upon reasonable request. Source data are included in this published article (and its supplementary information files). RNA-seq data have been deposited in public database.

## Methods

### Mice

C57BL/6 mice (Jax 664), CD45.1 (Jax 002014), μMT (Jax 2288) mice were originally from the Jackson Laboratory. Mb1-cre (*Cd79a*^cre/+^) mice were a gift from M. Reth. *Aicda*^CreERT^^2/+^ mice were a gift from JC Weill. *S1pr2*-cre^ERT^^2^ mice were a gift from T. Kurosaki. B1-8^hi^ immunoglobulin heavy chain knock-in mice were a gift from M. Nussenzweig. *Il9*^-/-^ mice were previously published ^9^. *Zbtb18*^fl/fl^ mice and *Zbtb18*^Flag-in/Flag-in^ knock-in mice were created on the B6 background using CRISPR/Cas9 technology by Biocytogen Ltd. To create the conditional *Zbtb18* knockout allele, two loxP sites were inserted into introns that flank exon 2, which contains the entire coding sequence of *Zbtb18*. To create a knock-in allele for expressing 3×Flag tagged Zbtb18, a GGS linker followed by a 3×Flag tag was inserted immediately before the stop codon for *Zbtb18* (Supplementary Fig. 1). Correct targeting was verified by sequencing and Southern blotting. All mice were maintained under specific pathogen-free conditions and used in accordance of governmental and Tsinghua guidelines for animal welfare.

### Generation of bone-marrow chimeras

A total of 2×10^6^ bone-marrow leukocytes containing 80% μMT cells and 20% *Il9*^-/-^ or *Il9*^+/+^ cells were transferred into the B6 recipient mice that were lethally irradiated by X-ray (2×5.5Gy). Chimeric animals were used 8 weeks after reconstitution.

### Immunization

To measure primary immune response, mice were immunized intraperitoneally with 100 μg NP-KLH (conjugate ratios:220, Biosearch Technologies) mixed with 1 μg lipopolysaccharide (LPS, Sigma) in alum (Thermo Scientific). To prepare OVA-primed mice for MBC transfer, mice were intraperitoneally immunized with 20 μg OVA mixed with 2 μg LPS in alum. To measure the recall response, mice were immunized intraperitoneally with 20 μg NP-OVA (Biosearch Technologies) in PBS. For sheep red blood cells (SRBC) immunization, each mouse was injected with 1 ml SRBC (Beijing Solarbio Science & Technology).

### Flow cytometry

Single-cell suspension of splenocytes was stained in MACS buffer (PBS supplemented with 1% FBS and 5 mM EDTA) containing Fc blocker (supernatant from 2.4G2 cell culture) for 20 min on ice before surface staining with primary antibodies in appropriate concentrations. Antibodies included PE-Cy7 anti-CD95 (Jo2), APC-Cy7 anti-CD19 (1D3), APC-Cy7 anti-CD45R (RA3-6B2), biotin anti-CD138 (281-2), Streptavidin PE (554061) (all from BD Biosciences); AF488 anti-GL7 (GL-7), eFluor450 anti-GL7 (GL-7), ef660 anti-GL7 (GL-7), biotin anti-GL7 (GL7), APC anti-Va2 TCR (B20.1) (all from eBiosciences); APC anti-CD86 (GL1), FITC anti-CD86 (GL1), PE anti-CXCR4 (L276F12), APC anti-CXCR4 (L276F12), FITC anti-CD4 (GK1.5), FITC anti-CD45.2 (104), AF700 anti-CD38 (90), BV510 anti-CD138 (281-2), PerCP Cy5.5 anti-IgD (11-26c), PE anti-73(catalog:127205), Streptavidin APC (catalog:405207), FITC anti-IgG (poly4053), APC anti-IgM (RMM-1) (all from Biolegend); NP-PE and NIP-biotin (both from Biosearch Technologies). Dead cells and doublets were excluded from analyses based on staining of 7-amino-actinomycin D (7-AAD) (Biotium) or Zombie yellow (catalog: 423103, Biolegend) and characteristics of forward and side scatter. Data were collected on a LSR II, or a Symphony A5, or an Aira III (BD Biosciences). Additional data were collected on an AURORA (Cytek). All data were analyzed with FlowJo software (TreeStar).

### Il-9 stimulation of GC B cells ex vivo

GC B cells were magnetically enriched with anti-GL7 antibody from B6 mice that were intraperitoneally immunized with 300 μl sheep red blood cells (SRBC, Beijing Solarbio Science & Technology). GC B cells were then FACS sorted into dark zone cells (CD19^+^GL7^+^FAS^+^CXCR4^+^CD86^-^) and light zone cells (CD19^+^GL7^+^FAS^+^CXCR4^-^CD86^+^) and cultured in the presence of 200 ng/ml Il-9 (PeproTech) in RPMI media for 3 h at 37°C. Total RNA was then extracted and reverse-transcribed into cDNA for quantitative RT-PCR.

### Quantitative RT-PCR

Total RNA was extracted by the Hipure Total RNA Micro KIT (Magen) and then reverse-transcribed into cDNA by 5×All-In-One RT Master Mix (Abm). For some experiments, cDNA libraries constructed with the Smart-seq2 protocol were used as templates. The EvaGreen Master Mix (Abm) was used to set up SYBR Green-based quantitative RT-PCR reactions following the manufacturer’s protocol. Samples were analyzed on a CFX Connect Real-Time system (Bio-RAD) and melting curves were used to validate correct amplification products. Expression levels of target genes were normalized to generic house-keeping gene *Actb* or the B-cell essential gene *Cd19,* which is expressed by naïve, activated, GC and memory B cells more uniformly. Primers used include the following: *Actb* sense 5′-GCTTCTTTGCAGCTCCTT, antisense 5′- TGCCAGATCTTCTCCATGT; *Cd19* sense 5’-AAATCCACGCATTCAAGTC, antisense 5’- TTCTCATAGCCACTCCCATCC; *Zbtb18* sense 5’- TGACTTTTCCCACGTCTTTAAGT, antisense 5’- AAACTCTGAGCTGGCATGGG; *S1pr2* Sense 5’ -CAACTCCGGGACATAGACCG, antisense 5’-CCAGCGTCTCCTTGGTGTAA; *Ccna2* sense 5’-TTGTAGGCACGGCTGCTATGCT, antisense 5’- GGTGCTCCATTCTCAGAACCTG; *Ccnb1* sense 5’- AGAGGTGGAACTTGCTGAGCCT, antisense 5’-<colcnt=3> GCACATCCAGATGTTTCCATCGG; *Ccnb2* sense 5’- GCACTACCATCCTTCTCAGGTG, antisense 5’-TGTGCTGCATGACTTCCAGGAC; *Cdk1* sense 5’-CATGGACCTCAAGAAGTACCTGG, antisense 5’- CAAGTCTCTGTGAAGAACTCGCC; *Cdk2* sense 5’- TCATGGATGCCTCTGCTCTCAC, antisense 5’- TGAAGGACACGGTGAGAATGGC; *Casp3* sense 5’- GGAGTCTGACTGGAAAGCCGAA, antisense 5’- CTTCTGGCAAGCCATCTCCTCA; *Bid* sense 5’- CCACAACATTGCCAGACATCTCG, antisense 5’- TCACCTCATCAAGGGCTTTGGC.

### Western blotting

Desired B cell subsets were sort-purified from *Zbtb18*^Flag-in/Flag-in^ mice 14 days after NP-KLH immunization. Cells were lysed in RIPA buffer containing a proteinase inhibitor cocktail. Whole-cell lysates were electrophoresed on SDS gels and transferred to nitrocellulose membrane. Samples were incubated with an anti-Flag (Catalog: 14791, Cell Signaling Technology) or anti-GAPDH antibody (Catalog: BE3407, EASYBIO) overnight at 4°C, and targets were revealed by staining with HRP Anti-rabbit secondary antibodies for 2 h at the room temperature. Images were acquired with Amersham Imager 600.

### Transduction of B cells

Retrovirus was generated by transfecting the Plat-E packaging cell line with 2.5 μg DNA and 7.5 μl TransIT-X2 Reagent (Mirus, MIR 600). B cells were isolated and cultured in complate 1640 RPMI medium before transduction. For transduction, the virus mixture was supplemented with 4 ng/ml polybrene and pre-heated in 37°C. B cells were spin-infected with viral supernatant at 1500 g in 37°C for 2 hours on day 1 and day 2. Transduced B cells were sorted for the qRT-PCR analysis or transferred into the recipients.

### Calcium assay following BCR stimulation

B cells were isolated with anti-mouse CD19 microbeads (Catalog:130-097-144, Miltenyi Biotec) from *Cd79a*^cre/+^*Zbtb18*^fl/fl^ and *Cd79a*^cre/+^ mice. B cells were stained with 2 μg/ml Indo-I (Catalog:I1203, Invitrogen) in RMPI medium for 30 mins at 37 °C, washed with cold PBS, and further stained for appropriate surface markers for 30 mins on ice. After washes, cells were re-suspended in RPMI containing 1% FBS, warmed at 37 °C in a water bath for 5 mins. Indo bound/Indo free ratio baseline was recorded for 1 min before addition of pre-warmed anti-mouse IgM F(ab’)_2_ (Jackson ImmunoResearch) to a final concentration of 2 or 10 μg/ml. Calcium flux was continuously recorded for a total of 5 mins.

### B-cell proliferation in vitro

B cells were isolated with anti-mouse CD19 microbeads from *Cd79a*^cre/+^*Zbtb18*^fl/fl^ and *Cd79a*^cre/+^ mice. B cells were stained with CellTrace^TM^ Violet (1:1000 in PBS; Catalog:34557, Invitrogen) for 30 mins at 37 °C. After washes, B cells were cultured in the presence of 1 μg/ml LPS, or 2 μg/ml anti-mouse IgM F(ab’)_2_ (Jackson ImmunoResearch), or 5 μg/ml anti-mouse CD40 (clone FGK4.5/ FGK45, BioXcell) for 72 hours before the dye dilution was analyzed by flow cytometry.

### Transwell assay

*Cd79a*^cre/+^*Zbtb18*^fl/fl^ and *Cd79a*^cre/+^ mice that were intraperitoneally immunized with 300 μl sheep red blood cells (SRBC, Beijing Solarbio Science & Technology), and GC B cells were enriched on day 9 by depleting all CD43^+^ and all IgD^+^ cells with anti-mouse CD43 microbeads (Catalog:130-097-148, Miltenyi Biotec) and biotinylated anti-mouse IgD/anti-biotin microbeads (Catalog:130-090-485, Miltenyi Biotec). Cells were pooled from 3 donor mice of each genotype in each experiment. B cells were suspended at 2×10^6^ cells per milliliter and rested in RPMI medium containing 1% low-lipid BSA (SRE0098, Sigma) at 37 °C for 1 h. Cells were then suspended in RPMI media containing 1% low-lipid BSA and 100 nM S1P (S9666, sigma) or 100 nM S1P plus 20 μM JET-013 (2392, Tocris). A total of 100 μl cell suspension as prepared above was added to the upper chamber in a transwell (pore size 5 μm, Corning Costar), and a total of 150 μl RPMI containing 0.3 μg/ml CXCL12 (Peprotech) was added to the lower chamber. Cells were allowed to transmigrate for 3 h at 37 °C in an incubator. CD38^+^IgD^-^Fas^+^GL7^+^ that had migrated to the bottom were enumerated by flow cytometry after surface staining. Separate wells were loaded with cell suspension without the transwell insert to serve as the input. A known number of carrier cells were added to each sample immediately before cytometry reading for cell counting.

### Recall assays of memory B cells

To recall polyclonal NP-specific MBCs, donor splenocytes were depleted of CD4^+^, CD8^+^, GL7^+^, CD138^+^ and IgD^+^ cells using a cocktail of biotinylated anti-CD4, anti-CD8, anti-GL7, anti-CD138, anti-IgD antibodies in combination of streptavidin Microbeads (Miltenyi Biotec). CD19^+^ cells were then enriched using CD19 Microbeads (Miltenyi Biotec). The resulting memory B cell-enriched cell preparation was FACS-analyzed, normalized for the abundance of NIP-binding B220^+^CD138^-^GL7^-^FAS^-^CD38^+^IgD^-^ MBCs, and transferred into CD45.1 recipients, which were primed with carrier protein OVA (Catalog:A5503, Sigma) 30 days before. After the transfer, recipients were immunized with NP-OVA and sacrificed 7 days later to enumerate GCs and PCs. To assay MBC recall using B1-8^hi^ cells, 5×10^5^ GFP-expressing B1-8^hi^ cells were adoptively transferred into OVA-primed *Il9*^+/+^ or *Il9*^-/-^ recipients that were subsequently immunized with NP-OVA. On day 7 post immunization, splenocytes from those immunized recipients were depleted of CD4^+^, CD8^+^, GL7^+^, CD138^+^ and IgD^+^ cells as described above. Resulting GFP^+^ NIP-binding MBCs were sort-purified, and 3000 MBCs were transferred into CD45.1 recipients that were primed with OVA 7 days earlier. These CD45.1 recipient mice were immunized with NP-OVA and sacrificed 7 days later to enumerate GFP^+^ B1-8^hi^ GCs and PCs.

### RNA-seq analysis of GC memory precursors

GC B cells were enriched from pooled splenocytes of 3 to 4 *Cd79a*^cre/+^*Zbtb18*^fl/fl^ or *Cd79a*^cre/+^ mice with anti-mouse GL7 (clone GL7, eBioscience) 28 days after NP-KLH immunization. Memory precursor cells (GL7^+^FAS^+^CD38^+^IgD^-^CD19^+^) were then sorted into 4 replicates of ∼200 cells per PCR tube containing the lysis buffer. Samples were reverse transcribed with SuperScript™ II Reverse Transcriptase (Catalog:18064022, Invitrogen) and the complement DNA library was constructed with TruePrep DNA Library Prep Kit for Illumina (Catalog:TD501, Vazyme), following the Smart-seq2 protocol. Amplified products were purified with VAHTS DNA Clean Beads (Vazyme) and quantified using Qubit and 2100 Bioanalyzer (Agilent). Quantitative RT-PCR was used for quality control. All libraries were sequenced on a HiSeq X Ten sequencer (Illumina). After pre-processed using FastQC (v0.11.9), raw sequences were aligned to the reference genome (GRCm38) using Hisat2 (version 2.2.1) and then sorted with Samtools (version 1.10). Reads then were counted in genes with the utilization of HTSeq-count (version 0.12.4). Genes that have a total read number of at least 1 were taken into downstream analysis. Differentially expressed genes (*P*_adj_<0.05) were calculated by the DESeq2 software in R. Ig genes were removed before calculation of adjusted *P* values to eliminate biases introduced by the abundantly expressed immunoglobulins. KEGG pathway-enrichment analysis were conducted with software GSEA 4.1.0.

### ATAC-seq

ATAC-seq were performed using TruePrep^TM^ DNA Library Prep Kit V2 for Illumina (Vazyme TD501). Briefly, 5×10^4^ naïve B cells or MBCs were sorted from *Cd79a*^cre/+^ mice and *Cd79a*^cre/+^*Zbtb18*^fl/fl^ mice, respectively. Cells were collected by gentle centrifugation (500x g, 5 minutes) at 4 °C, and then resuspended in lysis buffer (10 mM Tris.Cl, 10 mM NaCl, 3 mM MgCl_2_, 0.1% (v/v) Igepal CA-630) and incubated on ice for 10 min. Nuclei were collected and resuspended into 50 μl tagmentation mix, which contains 10 μl 5×TTBL (Vazyme), 5 μl TTE Mix V50 and 35 μl ddH_2_O. Samples were incubated at 37 °C for 30 min. After tagmentation, fragmented DNA was isolated by using VAHTS DNA Clean Beads. ATAC libraries were amplified for 12 PCR cycles using TruePrep^TM^ Index Kit V2 for Illumina (Vazyme #TD202). 0.55×VAHTSTM DNA Clean Beads were used to exclude the High–molecular weight DNA, followed by positive selection with 1×beads. High-quality DNA libraries were sequenced on the Illumina HiSeq2000.

### Luciferase assay

293T cells cultured in 48 well plate were co-transfected with GL3 empty vector or GL3 vector containing the wildtype *Ccna2*, *Ccnb1*,*Ccnb2*, *Cdk1*, *Cdk2*, *S1pr2*, *Casp3* and *Bid* promoter or mutated *S1pr2* promoter (313 ng/well), together with MSCV-GFP control vector or a vector expressing Zbtb18-3×Flag fusion protein (313 ng/well). RLTK vector (6.25 ng/well) was co-transfected as the internal control. After 48 h, cells were collected to measure the firefly luciferase and renilla luciferase luminescent activities using the Dual-Luciferase Reporter Assay kit (Catalog:E1910, Promega) with 96-well plates (Nunc MaxiSorp) and luminometer Glomax Multiplus (Promega), following the manufacturer’s protocol.

### ChIP PCR

ChIP was performed using the Hyperactive^TM^ In-Situ ChIP Library Prep Kit (Catalog:TD901-TD902, Vazyme). Briefly, one million GCMP cells were collected from *Zbtb18*^Flag-in/Flag-in^ mice and bound to the ConA beads. The cells were then suspended in 50 µl antibody buffer pre-mixed with an anti-Flag antibody (1:50; Catalog:14793, Cell signaling Technology) or with anti IgG control (Catalog:2729S, Cell signaling Technology) and incubated at the room temperature for 2 h, followed by incubation with the secondary antibody (Catalog:ab6701, abcam) and pG-Tn5 transposase. After extensive fragmentation, DNA was extracted with PCI (Phenol:Chloroform:Isoamyl alcohol), precipitated with ethanol, and suspended in 20 µl ddH_2_O. Samples were used for PCR amplification, and PCR products were purified with VAHTS DNA Clean Beads (Vazyme) and quantified using Qubit and 2100 Bioanalyzer (Agilent). The *Ccna2*, *Ccnb1*,*Ccnb2*, *Cdk1*, *Cdk2*, *S1pr2*, *Casp3* and *Bid* gDNA level was quantified by qPCR and *Actin* gDNA was used as internal control. The primer pairs were as follows: *Ccna2*: sense 5’- CCCATTGTTAGAAGGTTATCAA, antisense 5’- ATTCAACACATACAAGCCTGG; *Ccnb1*: sense 5’- GCAGTCGCTATTGGGGAGCTT, antisense 5’-<colcnt=4> TGCCTCCCAAGTGCTGGGATTA; *Ccnb2*: sense 5’- CCAAAGCTTCCCAAGAAAGAG, antisense 5’-TAAGTTCTCTCAAGTACCTGCT; *Cdk1*: sense 5’- TACACACAGAAAGGTAGCTGGA, antisense 5’CAACTGACTCTTGAGCTCTG; *Cdk2*: sense 5’-TTTCATGTGATGCTCAGCTGAGAC, antisense 5’- GGGTAATATCTTAAATGCAGTGATA; *S1pr2*: sense 5’ - ACATCTGGAATTCATTGCAA, antisense 5’-GTGGTTGGTTTTGGAGACTGTT; *Casp3*: sense 5’ -CTCTGAGTCCCCACAGTGAGGA, antisense 5’- TGCCGTTAATGTTGACTTCATC; *Bid*: sense 5’ - TGTGTGTTGCCTGCATATATGT, antisense 5’- TAGCACCCATATGGTGGCTGAC; *Actin*: sense 5’- CGTATTAGGTCCATCTTGAGAGTAC, antisense 5’-GCCATT GAGGCGTGATCGTAGC.

### Detection of active caspases ex vivo

CaspGLOW^TM^ Fluorescein Active Caspase Staining Kit (Catalog:88-7003, Invitrogen) was used according to the manufacturer’s protocol to measure active caspases in CD38^+^IgD^-^Fas^+^GL7^+^ memory precursor cells.

### Immunofluorescence

To examine expression patterns of Stat1 and Zbtb18, *Zbtb18*^Flag-in/Flag-in^ GCMP cells were treated with or without IL-9 for indicated amount of time and then fixed with 1% paraformaldehyde for 30 min on ice, washed in PBS for three times. Stat1 was revealed by staining with an anti-Stat1 antibody (Abcam Cat# ab239360) for 2 h, followed by staining with a donkey anti-rabbit secondary antibody for 1 h. An APC-conjugated anti-Flag antibody was used to stain Ztbtb18. Cells were examined with a Zeiss LSM780 upright confocal microscope with 40× air lens. Image data were analyzed in ZEN2 software (Zeiss). To examine distribution patterns of GCMP cells, splenic tissue collected from NP-KLH immunized mice on day 14 were fixed with 4% paraformaldehyde for 6 h at 4 °C and then dehydrated in 30% sucrose solution overnight at 4 °C. The GC-specific marker Ephrin B1 was revealed by overnight staining with purified goat anti-Ephrin B1 (catalog: AF473, R&D), followed by staining with AF647. IgD was stained by APC anti-IgD (clone:11-26c, 2123760, eBioscience) and CD38 was stained by ef450 anti-CD38 (catalog:E16165-104, eBioscience) for 2 h. Frozen slides were mounted with the ProlongGold Antifade reagent (Invitrogen) and examined with an Zeiss LSM780 upright confocal microscope.

### Statistics

Statistics and graphing were done in Prism (Graphpad). Unless indicated otherwise, two-tailed unpaired Student’s *t* test was used to compare end-point means of different groups.

---

**Supplementary Figure 1.**
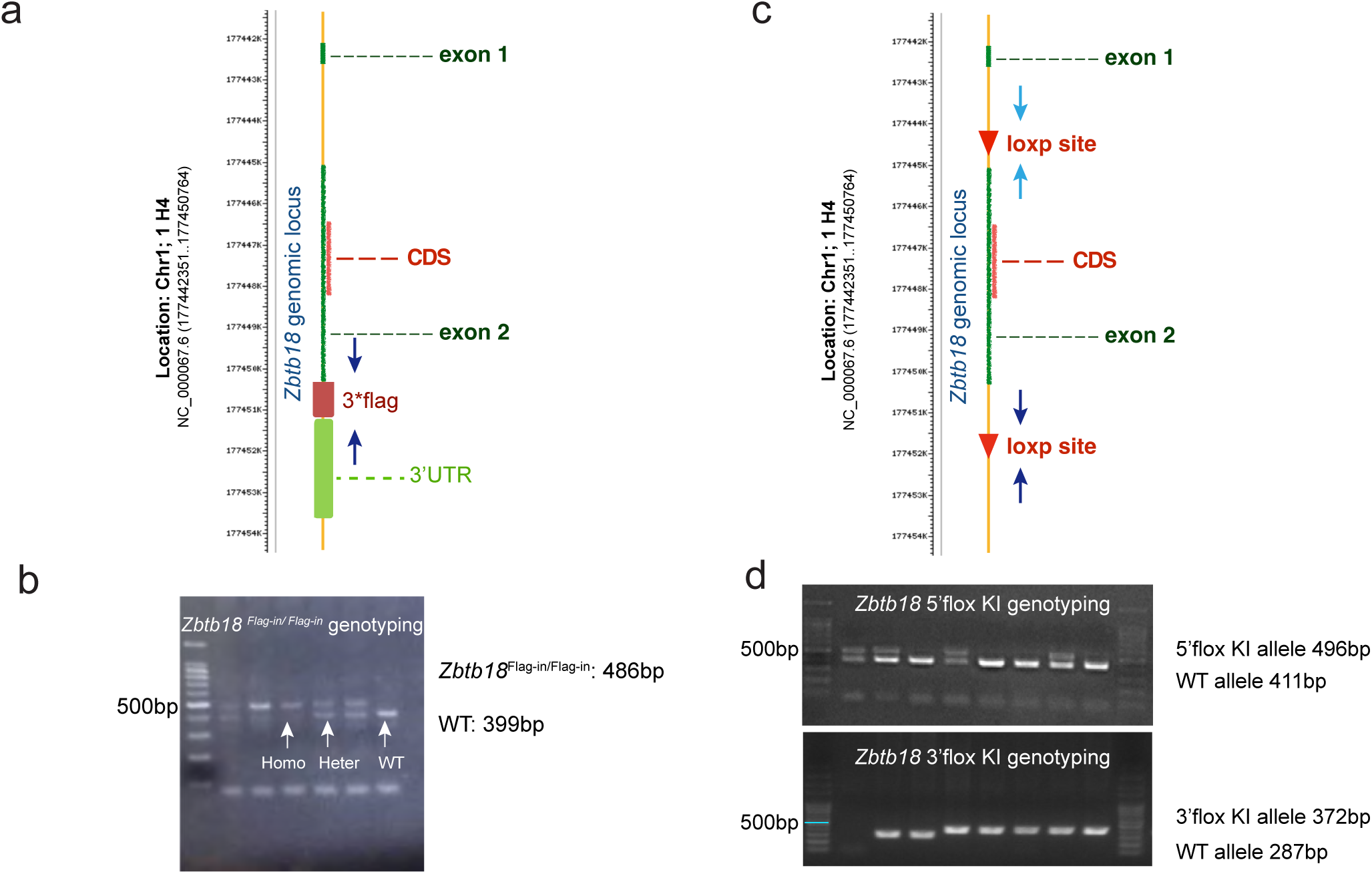
Construction of Zbtb18 conditional knockout and Flag-tagged allele. **a,** A schematic diagram of the *Zbtb18*^Flag-in^ allele. Positions of genotyping primers are indicated with arrows. **b,** Examples of genotyping results. **c,** A schematic diagram of the floxed *Zbtb18* allele. LoxP sites are indicated by arrowheads. Positions of two sets of genotyping primers are indicated with arrows. **d,** Examples of genotyping results.

**Supplementary Figure 2.**
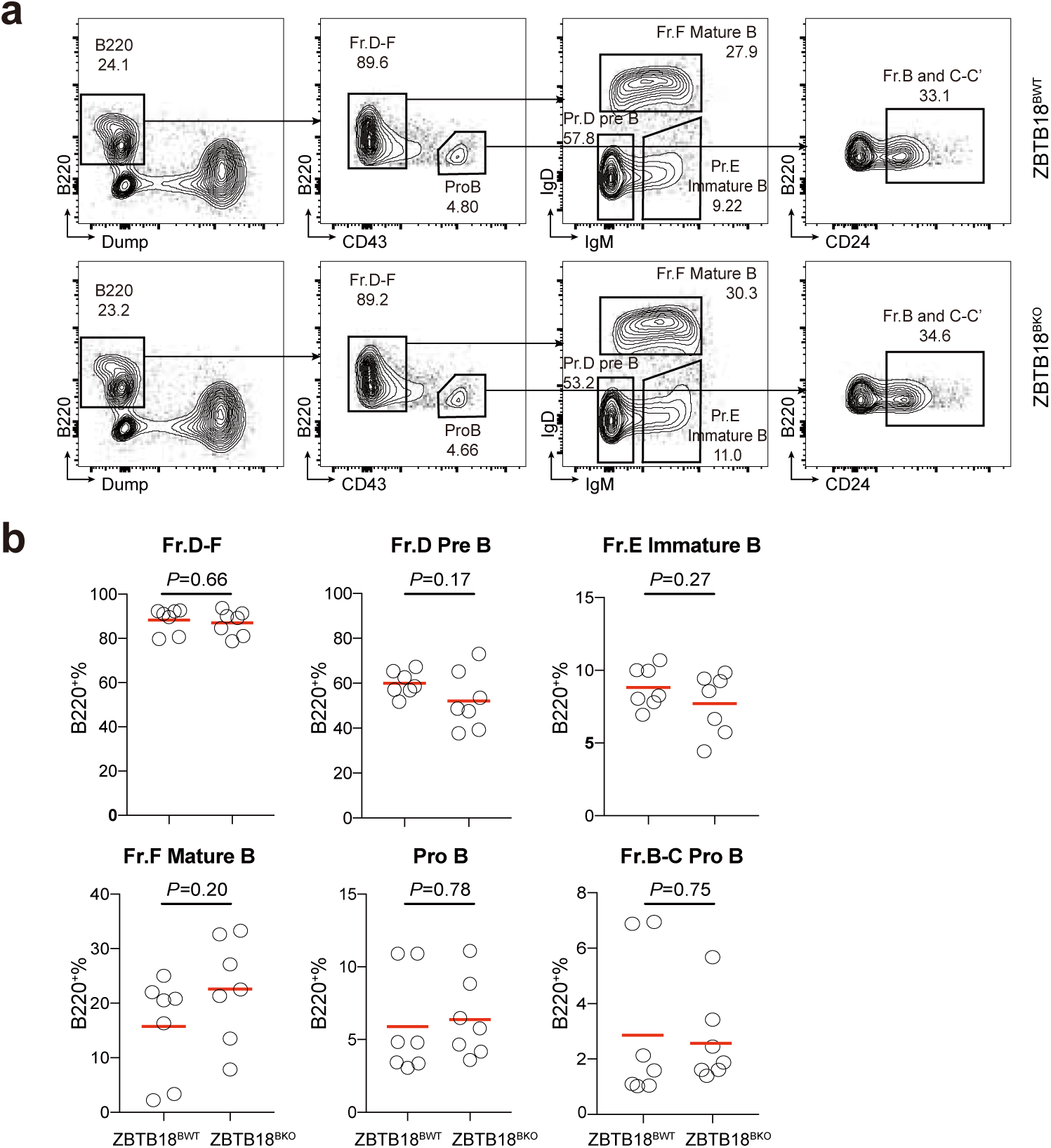
Bone-marrow B cell development in *Cd79a*^cre/+^ and *Zbtb18*^fl/fl^*Cd79a*^cre/+^ mice. **a,** Gating of bone-marrow B cells in ZBTB18^BWT^ and ZBTB18^BKO^ mice. **b,** Summary data of different developmental stages of B cells as gated in (**a**). Each symbol represents one mouse, and lines denote means. Data are pooled from 2 independent experiments. *P* values from *t* tests.

**Supplementary Figure 3.**
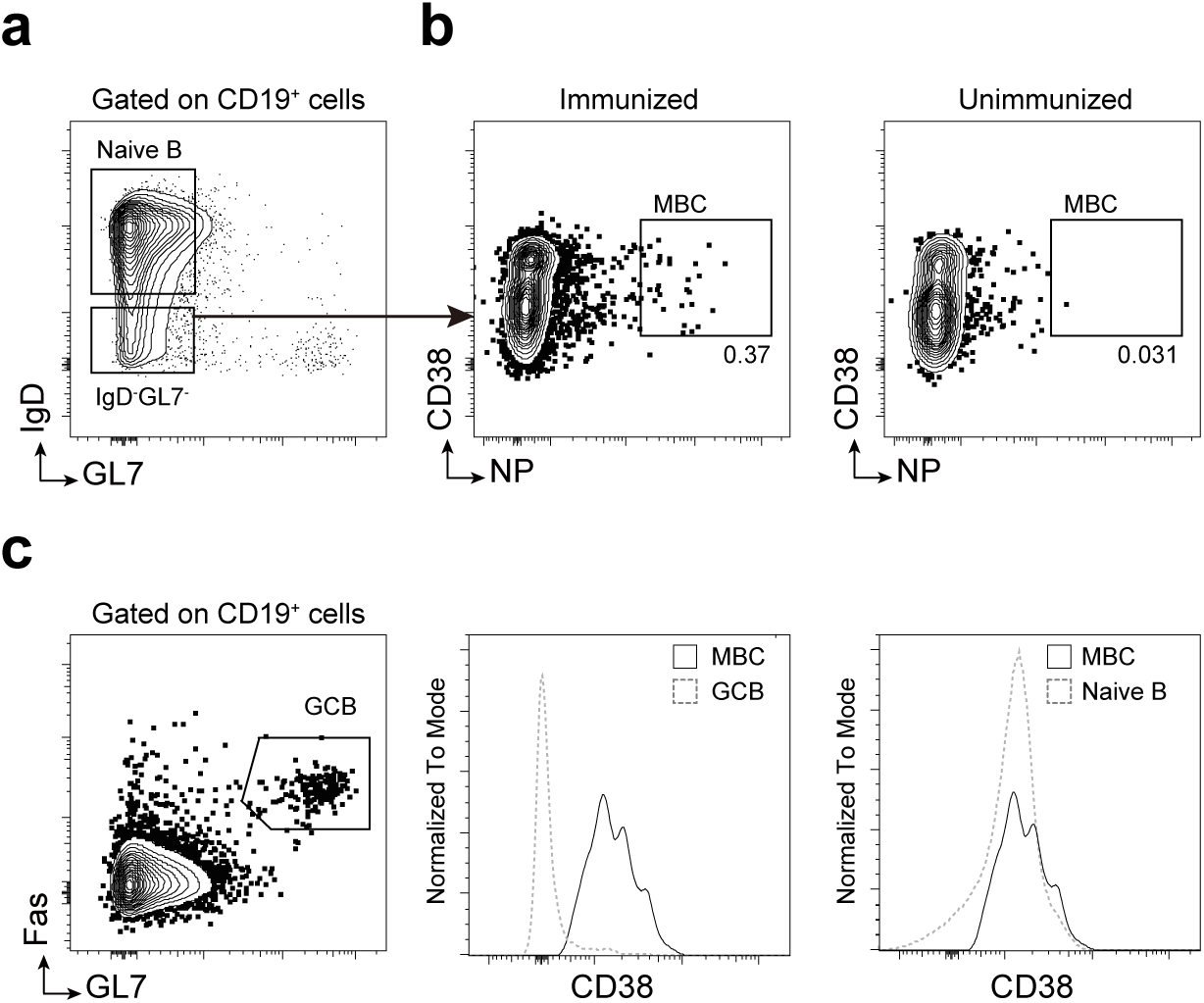
Gating of CD38^+^ NP-binding MBC cells in B6 mice. **a,** MBCs are gated from the IgD^-^GL7^-^CD19^+^ splenocytes. **b,** The NP-binding gate for IgD^-^GL7^-^CD19^+^ splenocytes from immunized mice (Left) is set according to that from unimmunized mice (Right). **c,** The CD38^+^ threshold for MBCs is set above the bulk GC B cells (as gated in the left panel) and comparable to the bulk naïve B cells (as gated in **a**).

**Supplementary Figure 4.**
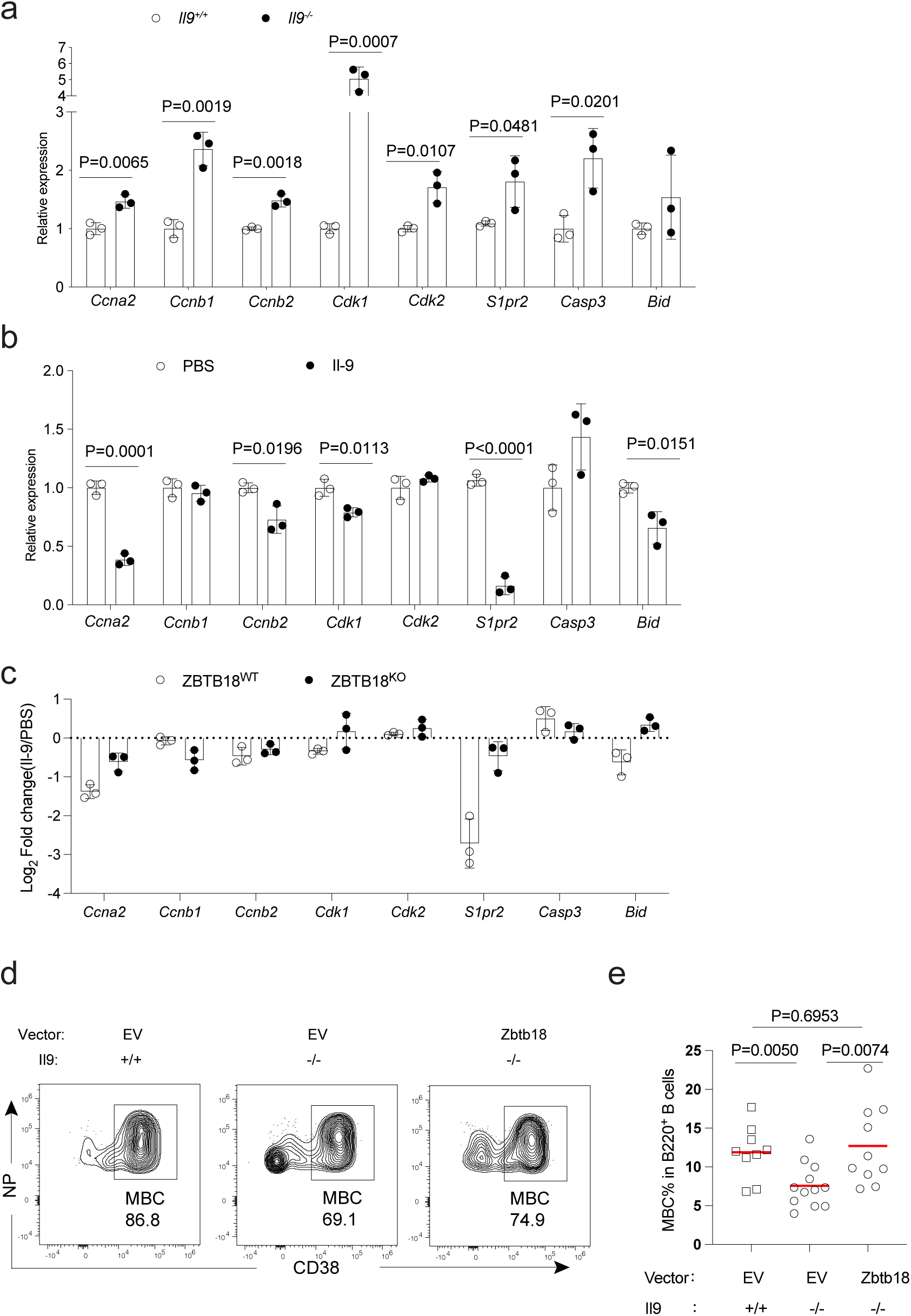
Il-9-Zbtb18 axis regulates MBCs. **a,** Relative mRNA expression of indicated Zbtb18 target genes in *Il9*^+/+^ and *Il9*^-/-^ GCMP cells. **b,** Relative mRNA expression of indicated Zbtb18 target genes in GCMP cells after 3h treatment with Il-9 *ex vivo*. Data are pooled from 3 independent experiments, *P* values by *t* tests. **c,** Effects of Il-9 treatment on indicated Zbtb18 target genes in GCMP cells sufficient or deficient in Zbtb18. Data presented as fold change between Il-9 treatment and the PBS control, pooled from 3 independent experiments. **d**, Representative FACS profiles of MBCs generated from vector-(EV) or Zbtb18-transduced B1-8^hi^ cells activated in wildtype or *Il9*^-/-^ mice 7 days after NP-KLH immunization. **e**, Fractional abundance of MBCs. *P* values from *t* tests.

## References

1. Weisel, F., and Shlomchik, M. (2017). Memory B Cells of Mice and Humans. Annual Review of Immunology 35, 255–284. 10.1146/annurev-immunol-041015-055531.

2. Liu, Y., Mason, D.Y., Johnson, G.D., Abbot, S., Gregory, C.D., Hardie, D.L., Gordon, J., and MacLennan, I.C.M. (1991). Germinal center cells express bcl-2 protein after activation by signals which prevent their entry into apoptosis. Eur. J. Immunol. 21, 1905–1910. 10.1002/eji.1830210819.

3. Liu, Y.-J., Joshua, D.E., Williams, G.T., Smith, C.A., Gordon, J., and MacLennan, I.C.M. (1989). Mechanism of antigen-driven selection in germinal centres. Nature 342, 929–931. 10.1038/342929a0.

4. Mayer, C.T., Gazumyan, A., Kara, E.E., Gitlin, A.D., Golijanin, J., Viant, C., Pai, J., Oliveira, T.Y., Wang, Q., Escolano, A., et al. (2017). The microanatomic segregation of selection by apoptosis in the germinal center. Science 358. 10.1126/science.aao2602.

5. Shinnakasu, R., Inoue, T., Kometani, K., Moriyama, S., Adachi, Y., Nakayama, M., Takahashi, Y., Fukuyama, H., Okada, T., and Kurosaki, T. (2016). Regulated selection of germinal-center cells into the memory B cell compartment. Nature Immunology. 10.1038/ni.3460.

6. Weisel, F.J., Zuccarino-Catania, G.V., Chikina, M., and Shlomchik, M.J. (2016). A Temporal Switch in the Germinal Center Determines Differential Output of Memory B and Plasma Cells. Immunity 44, 116–130. 10.1016/j.immuni.2015.12.004.

7. Wong, R., Belk, J.A., Govero, J., Uhrlaub, J.L., Reinartz, D., Zhao, H., Errico, J.M., D’Souza, L., Ripperger, T.J., Nikolich-Zugich, J., et al. (2020). Affinity-Restricted Memory B Cells Dominate Recall Responses to Heterologous Flaviviruses. Immunity. 10.1016/j.immuni.2020.09.001.

8. Mesin, L., Schiepers, A., Ersching, J., Barbulescu, A., Cavazzoni, C.B., Angelini, A., Okada, T., Kurosaki, T., and Victora, G.D. (2020). Restricted Clonality and Limited Germinal Center Reentry Characterize Memory B Cell Reactivation by Boosting. Cell 180, 92–106.e11. 10.1016/j.cell.2019.11.032.

9. Wang, Y., Shi, J., Yan, J., Xiao, Z., Hou, X., Lu, P., Hou, S., Mao, T., Liu, W., Ma, Y., et al. (2017). Germinal-center development of memory B cells driven by IL-9 from follicular helper T cells. Nature immunology 18, 921–930. 10.1038/ni.3788.

10. Laidlaw, B.J., Duan, L., Xu, Y., Vazquez, S.E., and Cyster, J.G. (2020). The transcription factor Hhex cooperates with the corepressor Tle3 to promote memory B cell development. Nat Immunol 21, 1082–1093. 10.1038/s41590-020-0713-6.

11. Laidlaw, B.J., Schmidt, T.H., Green, J.A., Allen, C.D.C., Okada, T., and Cyster, J.G. (2017). The Eph-related tyrosine kinase ligand Ephrin-B1 marks germinal center and memory precursor B cells. J Exp Medicine 214, 639–649. 10.1084/jem.20161461.

12. Suan, D., Kräutler, N.J., Maag, J.L.V., Butt, D., Bourne, K., Hermes, J.R., Avery, D.T., Young, C., Statham, A., Elliott, M., et al. (2017). CCR6 Defines Memory B Cell Precursors in Mouse and Human Germinal Centers, Revealing Light-Zone Location and Predominant Low Antigen Affinity. Immunity 47, 1142–1153.e4. 10.1016/j.immuni.2017.11.022.

13. Takatsuka, S., Yamada, H., Haniuda, K., Saruwatari, H., Ichihashi, M., Renauld, J.-C., and Kitamura, D. (2018). IL-9 receptor signaling in memory B cells regulates humoral recall responses. Nature Immunology 19, 1025–1034. 10.1038/s41590-018-0177-0.

14. Kaji, T., Ishige, A., Hikida, M., Taka, J., Hijikata, A., Kubo, M., Nagashima, T., Takahashi, Y., Kurosaki, T., Okada, M., et al. (2012). Distinct cellular pathways select germline-encoded and somatically mutated antibodies into immunological memory. J Exp Med 209, 2079–2097. 10.1084/jem.20120127.

15. Toyama, H., Okada, S., Hatano, M., Takahashi, Y., Takeda, N., Ichii, H., Takemori, T., Kuroda, Y., and Tokuhisa, T. (2002). Memory B Cells without Somatic Hypermutation Are Generated from Bcl6-Deficient B Cells. Immunity 17, 329–339. 10.1016/s1074-7613(02)00387-4.

16. Tarlinton, D., and Good-Jacobson, K. (2013). Diversity Among Memory B Cells: Origin, Consequences, and Utility. Science 341, 1205–1211. 10.1126/science.1241146.

17. Taylor, J.J., Pape, K.A., and Jenkins, M.K. (2012). A germinal center–independent pathway generates unswitched memory B cells early in the primary response. J Exp Med 209, 597–606. 10.1084/jem.20111696.

18. Anderson, S.M., Tomayko, M.M., Ahuja, A., Haberman, A.M., and Shlomchik, M.J. (2007). New markers for murine memory B cells that define mutated and unmutated subsets. J Exp Medicine 204, 2103–2114. 10.1084/jem.20062571.

19. Aoki, K., Meng, G., Suzuki, K., Takashi, T., Kameoka, Y., Nakahara, K., Ishida, R., and Kasai, M. (1998). RP58 associates with condensed chromatin and mediates a sequence-specific transcriptional repression. The Journal of biological chemistry 273, 26698–26704. 10.1074/jbc.273.41.26698.

20. Ohtaka-Maruyama, C., Hirai, S., Miwa, A., Heng, J.I., Shitara, H., Ishii, R., Taya, C., Kawano, H., Kasai, M., Nakajima, K., et al. (2013). RP58 regulates the multipolar-bipolar transition of newborn neurons in the developing cerebral cortex. Cell reports 3, 458–471. 10.1016/j.celrep.2013.01.012.

21. Clément, O., Hemming, I., Gladwyn-Ng, I., Qu, Z., Li, S., Piper, M., and Heng, J. (2017). Rp58 and p27kip1 coordinate cell cycle exit and neuronal migration within the embryonic mouse cerebral cortex. Neural Dev 12, 8. 10.1186/s13064-017-0084-3.

22. Okado, H., Ohtaka-Maruyama, C., Sugitani, Y., Fukuda, Y., Ishida, R., Hirai, S., Miwa, A., Takahashi, A., Aoki, K., Mochida, K., et al. (2009). The transcriptional repressor RP58 is crucial for cell-division patterning and neuronal survival in the developing cortex. Dev Biol 331, 140–151. 10.1016/j.ydbio.2009.04.030.

23. Fuks, F., Burgers, W.A., Godin, N., Kasai, M., and Kouzarides, T. (2001). Dnmt3a binds deacetylases and is recruited by a sequence-specific repressor to silence transcription. Embo J 20, 2536–2544. 10.1093/emboj/20.10.2536.

24. Green, J.A., and Cyster, J.G. (2012). S1PR2 links germinal center confinement and growth regulation. Immunological reviews 247, 36–51. 10.1111/j.1600-065X.2012.01114.x.

25. Xie, B., Khoyratty, T.E., Abu-Shah, E., Cespedes, P.F., MacLean, A.J., Pirgova, G., Hu, Z., Ahmed, A.A., Dustin, M.L., Udalova, I.A., et al. (2021). The Zinc Finger Protein Zbtb18 Represses Expression of Class I Phosphatidylinositol 3-Kinase Subunits and Inhibits Plasma Cell Differentiation. J Immunol 206, ji2000367. 10.4049/jimmunol.2000367.

26. Ersching, J., Efeyan, A., Mesin, L., Jacobsen, J.T., Pasqual, G., Grabiner, B.C., Dominguez-Sola, D., Sabatini, D.M., and Victora, G.D. (2017). Germinal Center Selection and Affinity Maturation Require Dynamic Regulation of mTORC1 Kinase. Immunity 46, 1045–1058.e6. 10.1016/j.immuni.2017.06.005.

